# A synaptic locus of song learning

**DOI:** 10.64898/2026.01.20.699515

**Authors:** Drew C Schreiner, Samuel Brudner, Amanda Li, John Pearson, Richard Mooney

## Abstract

Learning by imitation is the foundation for verbal and musical expression, but its underlying neural basis remains obscure. A juvenile male zebra finch imitates the multisyllabic song of an adult tutor in a process that depends on a song-specialized cortico-basal ganglia circuit, affording a powerful system to identify the synaptic substrates of imitative motor learning. Plasticity at a particular set of cortico-basal ganglia synapses is hypothesized to drive rapid learning-related changes in song before these changes are subsequently consolidated in downstream circuits. Nevertheless, this hypothesis is untested and the synaptic locus where learning initially occurs is unknown. By combining a computational framework to quantify song learning with synapse-specific optogenetic and chemogenetic manipulations within and directly downstream of the cortico-basal ganglia circuit, we identified the specific cortico-basal ganglia synapses that drive the acquisition and expression of rapid vocal changes during juvenile song learning and characterized the hours-long timescale over which these changes consolidate. Furthermore, transiently augmenting postsynaptic activity in the basal ganglia briefly accelerates learning rates and persistently alters song, demonstrating a direct link between basal ganglia activity and rapid learning. These results localize the specific cortico-basal ganglia synapses that enable a juvenile songbird to learn to sing and reveal the circuit logic and behavioral timescales of this imitative learning paradigm.

## Introduction

A juvenile male zebra finch memorizes the multisyllabic song motif of an adult male tutor and then engages in a prolonged (∼1 month) period of hearing-dependent vocal practice, known as sensorimotor learning, to imitate this memorized model^1–5^ (Fig. 1a). A song-specialized cortico-basal ganglia pathway is essential to this spontaneous form of sensorimotor learning^6–9^, affording the potential to identify the neural substrates that enable song imitation (Fig. 1b). An influential model is that two distinct cortical inputs into a song-specialized region of the basal ganglia (Area X, referred to here as the sBG) convey separate signals related to song timing and variability, the coincident activity of which generates a synaptic eligibility trace in the spiny neurons that make up >95% of sBG neurons^10–15^. In this model^15^, the initial acquisition and expression of sensorimotor learning depend on a dopamine-mediated reinforcement signal acting on the eligibility trace to selectively strengthen the time-encoding cortico-sBG spiny neuron synapses whose singing-related activity corresponds with song variants more like the tutor (Fig. 1c). This synaptic change alters sBG song premotor activity to bias the juvenile’s song towards the tutor model, an effect that is then consolidated over a longer timescale in motor regions downstream of the cortico-sBG pathway^16,17^.

**Figure 1.**
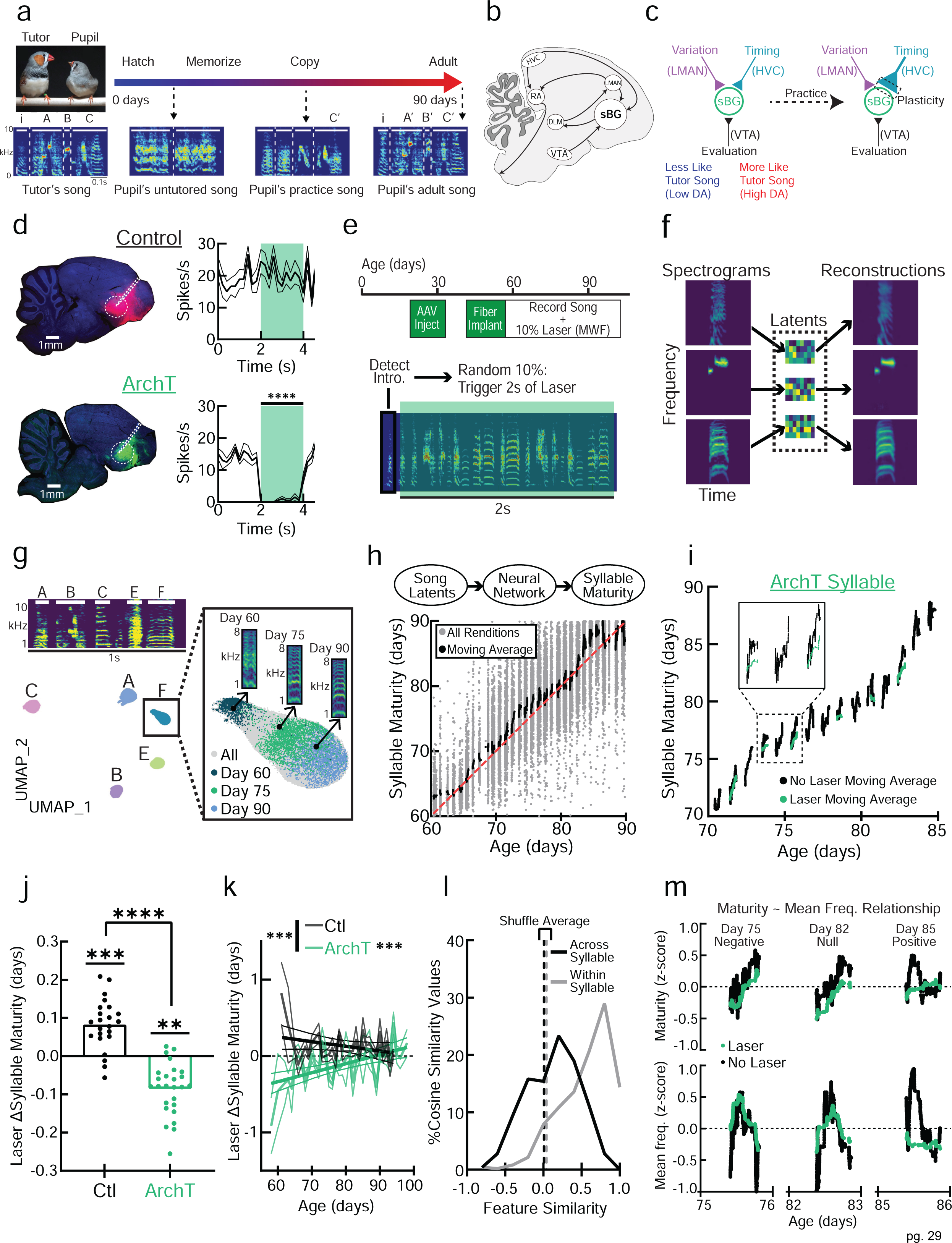
sBG activity is necessary for the expression of recent learning. (**a**) Schematic of juvenile song learning showing song spectrograms from the tutor and the pupil across learning, adapted from^2^. i = introductory note, A-C = syllables. (**b**) Sagittal diagram of the male zebra finch song system, including cortical nuclei HVC, LMAN, RA, song-specialized basal ganglia (sBG), DLM thalamus, and VTA. (**c**) Hypothesized model of song learning at HVC-sBG spiny neuron synapses^15^. (**d**) (Left) Representative histology for Control tdTomato (red) and ArchT-eGFP expression (green) with Neurotrace (blue). White circle = the sBG, dashed lines = optical fiber. (Right) Peri-event time histograms of sBG activity aligned to laser onset (green shading = laser) (ArchT: t-test t_19_ = 16.5, p < 0.0001). (**e**) Experimental timeline and schematic of laser administration. (**f**) Variational autoencoders (VAE) compress syllable spectrograms into a 32-dimensional latent embedding sufficient to decode an image closely matching the original spectrogram. (**g**) Sample song spectrogram (top) and UMAP representation (bottom) of VAE latent embeddings of the same song. (Right) Progression of syllable “F” from days 60-90. (**h**) (Top) Artificial Neural Network (ANN) trained to predict age of production based on the VAE’s latent embeddings for each syllable. (Bottom) Example ANN predicted Syllable Maturity vs. actual age for syllable “F”. (**i**) Moving average (200 renditions) of Syllable Maturity with/without laser for example ArchT syllable “F”. (**j**) Change (Δ) in Syllable Maturity when laser is on as predicted by a mixed effects model. Bar = fixed effect, points = estimated effects per syllable. Laser effects significantly differed between Ctl and ArchT (t_3.56e6_ = -4.98, p = 6.24e-7) and were significant in both Ctl (t_3.56e6_ = 3.57, p = 0.00072) and ArchT (t_3.56e6_ = -3.48, p = 0.0010). (**k**) Laser-induced change in Syllable Maturity grouped by day. Solid lines are a linear regression relating laser-induced maturity changes to age (ArchT: F_1,36_ = 17.6, p = 0.0002, R^2^ = 0.329. ArchT slope differs from Ctl: F_1,61_ = 14.5, p = 0.0003). (**l**) Histogram of cosine similarity values relating the vector of Feature regression coefficients in an elastic net regression predicting Maturity from Features either Across Syllables or Within Syllable across days. Dotted lines indicate the shuffled mean near 0 (real mean Across = 0.12, Within = 0.56). (**m**) An example syllable for which mean frequency (Freq.) is negatively (left panels), null (center), or positively (right) related to maturity on different days (lines are moving average). While sBG suppression consistently reduces maturity (top row), it affects mean frequency only when frequency is aligned with maturity (bottom row). Lines in k show Mean ± SEM. ** = p < 0.01, *** = p < 0.001, **** = p < 0.0001. ns = not significant.

Although this model^15^ is the dominant theoretical description of song learning, it remains largely untested. Whether sBG premotor activity actually drives progressive changes in song during juvenile learning remains unknown. Additionally, which synapses change within the sBG to drive the initial expression of song learning and the timescales over which these rapid changes undergo consolidation is unclear. However, one prediction of this model is that transiently suppressing sBG activity, and specifically the time-encoding cortico-sBG synapses, during singing will prevent the expression of recently learned changes in song but not changes that have already been consolidated. A second prediction of this model is that the initial acquisition of learning is localized to the sBG and requires activity in both the time- and variability-encoding cortico-sBG inputs.

To directly test these predictions, we combined computational methods to quantify rendition-to-rendition vocal changes during juvenile song learning^18,19^ with closed-loop optogenetic suppression or chemogenetic manipulations of specific pre- and postsynaptic components of the cortico-sBG circuit, as well as in downstream cortical premotor-motor circuitry. This approach enabled us to identify and distinguish synaptic loci crucial to the initial acquisition and expression of sensorimotor learning, quantify the timescale over which these neural and vocal changes undergo consolidation, and explore how sBG activity influences the quality and rate of learning.

### sBG activity is necessary to express recently learned changes to song

To establish a causal link between sBG activity and the expression of song learning, we quantified how transiently suppressing sBG activity affected rapid, learning-related changes in juvenile song. We expressed the inhibitory opsin Archaerhodopsin^20^ (ArchT) or a control fluorophore (tdTomato) in the sBG of normally tutored juvenile male zebra finches using viral vectors (injected 20 - 35 days post hatch (dph), n = 3 birds using AAV2/1-CAG-ArchT-eGFP for pan-neuronal expression; n = 3 birds using AAV2/1-CaMKIIa-ArchT-eGFP for spiny neuron specific expression^21^; n = 4 birds using AAV2/1-CAG-tdTomato as a control; each bird was housed in their home cage from hatching with their mother, father (an adult male tutor), and siblings). Three weeks later (41 - 56 dph), we verified that sBG action potential activity was suppressed by laser administration in ArchT- but not tdTomato-expressing birds and then implanted tapered optical fibers bilaterally in the sBG (Fig. 1d; Extended Data Fig. 1a). Following implantation, birds were isolated from their tutors and individually housed in sound attenuating chambers, tethered to optical fibers, and their songs were recorded continuously until ∼90 dph, when sensorimotor learning typically ends. On alternating weekdays, we detected the introductory notes^22^ that occur at the beginning of song motifs and administered 2 s of laser to the sBG on a random 10% of song renditions, a period sufficient to encompass one to several song motifs (Fig 1e).

To quantify the juvenile song learning trajectory, we segmented the entire corpus of recorded song motifs into their component syllables (3.6 million syllable renditions from 10 birds). We then used a Variational Autoencoder (VAE) to compress high dimensional (128 x 128 pixels, ∼16K) spectrographic data into 32 latent dimensions that preserved information sufficient to accurately reconstruct the original spectrograms^18^ (Fig. 1f).

Two-dimensional UMAP projections of these latent embeddings revealed that individual syllables were well-clustered and distinct from one another (Fig. 1g, left). Qualitatively, labeling syllable renditions by their age of production revealed that syllable representations in this latent space occupied different locations as a function of the age of production, corresponding to maturation of the underlying spectrogram (Fig. 1g, right). To quantify this relationship on a rendition-by-rendition basis, we trained an Artificial Neural Network (ANN) to predict the most likely age of production for each syllable rendition based on its location in this latent space^19^ (Fig. 1h). This VAE-ANN predicted age metric is hereafter referred to as “syllable maturity” or simply “maturity”.

The VAE-ANN method allowed us to assess how transiently suppressing sBG activity during singing affected syllable maturity. To quantify this, we built linear mixed effects models to predict how laser affected syllable maturity in both ArchT and tdTomato control birds, while also controlling for circadian and developmental contributions to maturity (see methods). Comparing laser-on to laser-off renditions in ArchT birds and in tdTomato fluorophore controls revealed that syllables were less mature when sBG activity was suppressed (Fig. 1i,j; ArchT: -2.05 ± 0.60 hours (Mean ± SEM), n = 26 ArchT syllables). This regressive effect of sBG suppression on syllable maturity occurred only on targeted syllables and was significantly different from laser shuffled data (Extended Data Fig. 1b,c). Finally, syllable maturity was significantly reduced in birds that expressed ArchT either pan-neuronally or only in spiny neurons in the sBG (Extended Data Fig. 1d). Thus, singing-related activity specifically in sBG spiny neurons is necessary for the expression of recently learned vocal changes during sensorimotor learning.

Motor learning studies in a variety of species, including zebra finches, indicate that recently learned skills are consolidated during sleep^23–25^. If sleep helps consolidate recently learned song changes downstream of the sBG, then the maturity of renditions produced in the morning just after sleep should be less affected by suppressing sBG activity. However, syllable maturity regressed to a similar extent regardless of when during the day sBG activity was suppressed (Extended Data Fig. 1e). We also examined the long-term consequences of intermittent sBG suppression. Final learning outcomes of syllables shared by the pupil and tutor were unaffected by this history of relatively infrequent (i.e. ∼10% of renditions) sBG suppression, underscoring that the protocol used here was able to successfully isolate the sBG’s role in expressing recently learned vocal changes (Extended Data Fig. 1f). Notably, the regressive effects on syllable maturity caused by suppressing sBG activity declined steadily over sensorimotor learning (Fig. 1k), consistent with various findings that highlight a greater role for cortico-sBG circuits in juvenile sensorimotor learning than in adult song production^6,7,9^.

Conceivably, suppressing sBG activity could regress syllable maturity by consistently altering certain acoustic features (e.g., pitch, amplitude) that are reliably associated with more mature syllables. However, individual acoustic features did not reliably relate to maturity measured across syllables and, even within syllables, which features predicted maturity could and did change from day to day (Fig. 1l, Extended Data Fig. 2a,b; see also^26^). Moreover, suppressing sBG activity did not reliably affect individual acoustic features across syllables or within the same syllable across days (Extended Data Fig. 2c,d). In fact, suppressing sBG activity most strongly affected individual acoustic features *if and when* those features were aligned with syllable maturity (Fig. 1m, Extended Data Fig. 2e,f). And while variability measured across all acoustic features decreased slightly when sBG activity was suppressed, variability decreased marginally more in fluorophore control birds, (Extended Data Fig. 1g) which exhibited no regression in syllable maturity (Fig. 1j). Lastly, suppressing sBG activity did not affect intrasyllabic variability, rendition-to-rendition variability of syllable maturity, syllable syntax, or intersyllabic gaps (Extended Data Fig. 2d,h-j). Therefore, in juvenile finches, singing-related sBG activity selectively modulates acoustic features of syllables that align with daily changes in syllable maturity, rather than simply modulating a fixed set of acoustic features or altering the order or timing of syllables.

### The time-encoding cortico-sBG synapse is necessary to express recent learning

The sBG receives singing-related excitatory input from the cortical nuclei HVC and LMAN, which are hypothesized to convey song timing and variability signals, respectively^15^. To further isolate a synaptic locus necessary for the expression of sensorimotor learning, we used viral vectors to express an inhibitory opsin (Parapinopsin, or PPO, a G_i/o_-coupled opsin)^27^ bilaterally in either HVC or LMAN axon terminals in the sBG (GFP was expressed in HVC-sBG terminals in an additional control group, n = 5 birds per group) (Fig. 2a). Three weeks later (41 - 56 dph), we verified that laser applied to the sBG reduced spontaneous action potential activity in PPO- but not GFP-expressing birds (Fig. 2a). We also verified that optogenetically suppressing HVC terminals using PPO could prevent sBG activity evoked by electrical stimulation in HVC (n = 2 birds, Extended Data Fig. 3a). Following bilateral implantation of optical fibers targeting the sBG, we used the same approach described previously for ArchT experiments (Fig. 1e) to suppress either HVC-sBG or LMAN-sBG terminal activity on a small subset (∼10%) of motifs produced across sensorimotor learning (5.6 million syllable renditions from 15 birds).

**Figure 2.**
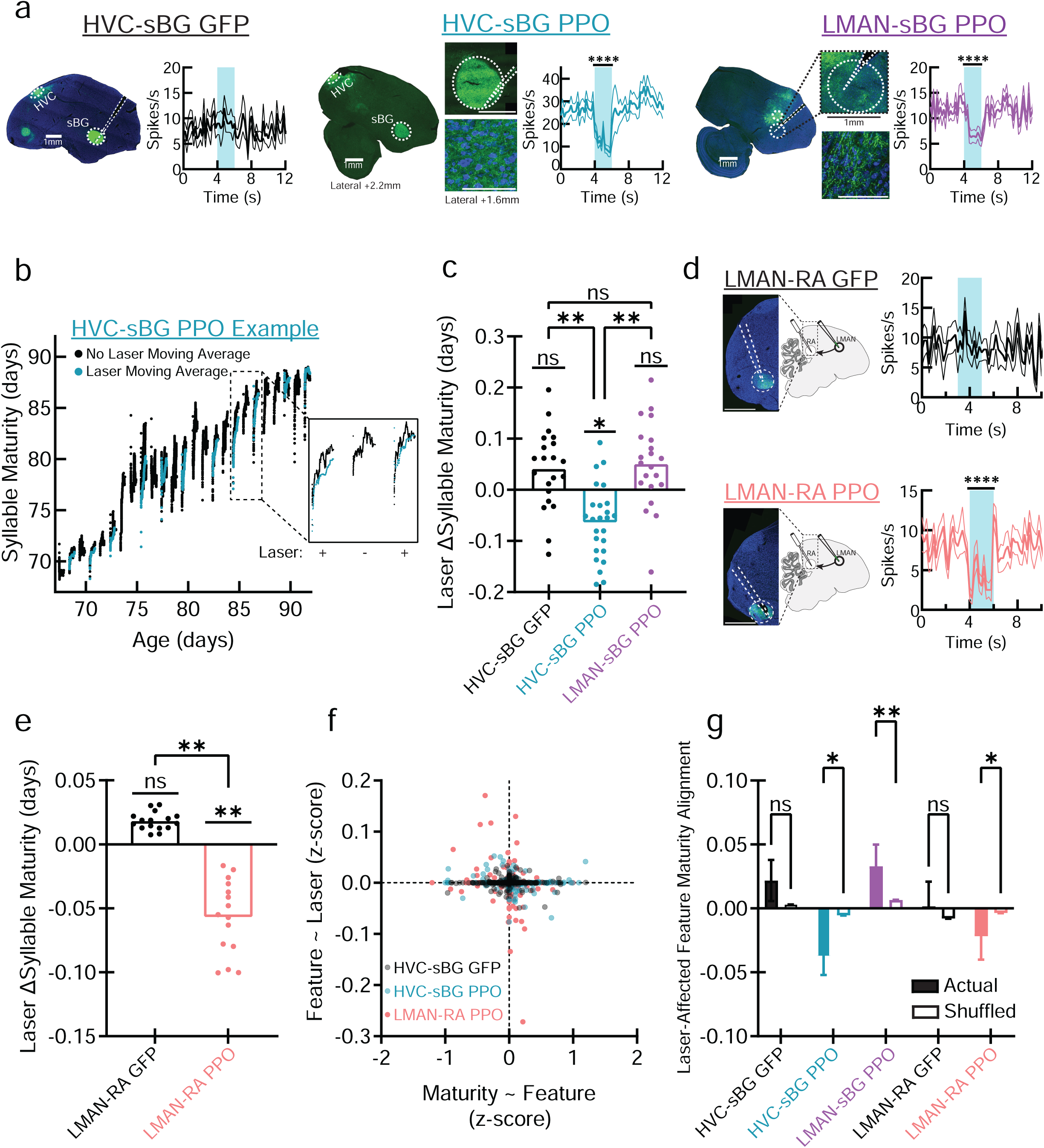
HVC-sBG and LMAN-RA terminals are necessary to express recent learning. (**a**) Representative histology and peri-event time histograms for HVC-sBG GFP (left), HVC-sBG PPO-mVenus (center), and LMAN-sBG PPO-mVenus (right) (green = mVenus label, blue = Neurotrace) and optical fiber tracks in the sBG (dashed white lines). HVC-sBG PPO shown at two lateral planes to capture HVC cell bodies (+2.2) and terminals in sBG (+1.6). Bottom row of HVC-sBG PPO and LMAN-sBG PPO show terminal fields at high magnification (scale bars = 100 µm). Example peri-event time histograms aligned to laser onset (blue shading) show robust suppression of sBG action potential activity in HVC-sBG PPO (unpaired t-test 2s before vs. 2s during laser: t_19_ = 17.9, p < 0.0001) and LMAN-sBG PPO (t_19_ = 6.34, p < 0.0001). (**b**) Syllable Maturity for example HVC-sBG PPO syllable. Laser +/- indicates days with/without laser. (**c**) Change (Δ) in Syllable Maturity when Laser is on relative to off as predicted by a mixed effects model. Bar = fixed effect, points = estimated effects per syllable. Corrected Laser effects in HVC-sBG PPO significantly differed from both HVC-sBG GFP (t_5.63e6_ = -3.12, p = 0.005), and LMAN-sBG PPO (t_5.63e6_ = -2.78, p = 0.0073). Post hoc corrected tests found a significant Laser effect only in HVC-sBG PPO. (t_5.63e6_ = - 2.64, p = 0.025). (**d**) Representative histology and peri-event time histograms for LMAN-RA GFP and LMAN-RA PPO show robust RA suppression (unpaired t-test: t_19_ = 6.18, p < 0.0001). (**e**) As in (**c**), but for the LMAN-RA experiment. Laser effects significantly differed between LMAN-RA GFP and LMAN-RA PPO (t_2.08e6_ = -3.06, p = 0.0031). Post hoc corrected tests found a significant Laser effect only in LMAN-RA PPO (t_2.08e6_ = - 3.16, p = 0.003). (**f**) Scatter plot relating elastic net coefficients for models predicting Maturity as a function of Features (Maturity ∼ Feature) vs. Feature ∼ Laser showing that, in HVC-sBG PPO and LMAN-RA PPO, when a feature predicts maturity (horizontal axis), Laser tends to push that feature in the opposite direction (vertical axis). Points represent the relationship between one syllable and one feature (e.g., Syllable A, Mean Frequency). (**g**) Alignment of laser-affected features with features that predict maturity. Cosine similarity of actual and shuffled data between the vector of acoustic features that predict maturity and the vector of laser effects on those same features. Positive numbers indicate that the effect of laser is aligned with maturity in feature space, while negative numbers indicate anti-alignment. HVC-sBG PPO and LMAN-RA PPO have more negative cosine similarities relative to shuffled data (HVC-sBG PPO: KS-test: D = 0.07 n_1_ = 344, n_2_ = 344 x 10000 shuffles, p = 0.045; LMAN-RA PPO: D = 0.10 n_1_ = 193, n_2_ = 193 x 10000 shuffles, p = 0.033), while the LMAN-sBG PPO group shows the opposite effect (D = 0.11 n_1_ = 258, n_2_ = 258 x 10000 shuffles, p = 0.005). * = p < 0.05, ** = p < 0.01, ns = not significant.

Comparing laser-on to laser-off renditions revealed that syllables were less mature when HVC-sBG, but not LMAN-sBG, terminals were suppressed (Fig. 2b,c HVC-sBG: -1.52 ± 0.58 hours, Mean ± SEM; n = 25 syllables). This regressive effect of HVC-sBG terminal suppression on maturity manifested only on laser-targeted renditions, was significant in comparison with laser shuffled data, and was not elicited by laser illumination of the sBG in GFP controls (Extended Data Fig. 3b,c). Variability measured across all acoustic features was unaffected by suppressing either HVC-sBG or LMAN-sBG terminals (Extended Data Fig. 3d). Along with the spiny neuron-specific sBG suppression experiments (Extended Data Fig. 1d), these observations indicate that the rapid expression of vocal changes during sensorimotor learning involves HVC axon terminal synapses on sBG spiny neurons^15^.

Prior studies showed that the expression of a simpler form of externally reinforced vocal learning in adult finches (“pitch learning”) depends on activity conveyed by the output of the cortico-sBG circuitry, namely the axonal projections from LMAN to the cortical song premotor nucleus RA^28^. To test whether a similar mechanism underlies the expression of juvenile sensorimotor learning, we explored how syllable maturity was affected by transient optogenetic suppression of LMAN-RA terminals during juvenile learning (n = 4 LMAN-RA PPO and n = 4 LMAN-RA GFP control birds; 2.1 million syllable renditions, Fig. 2d). Prior to behavioral experiments, we verified that spontaneous action potential activity in RA was suppressed by laser administration in PPO- but not GFP-expressing birds (Fig. 2d). Indeed, PPO-mediated suppression in RA was as efficacious as the cortico-sBG PPO manipulations (Extended Data Fig. 3e,f). Comparing laser-on to laser-off renditions revealed that syllables were less mature in LMAN-RA PPO but not GFP control birds (Fig 2e, LMAN-RA PPO: -1.36 ± 0.43 hours, Mean ± SEM; n = 15 syllables). This regressive effect of LMAN-RA terminal suppression on syllable maturity manifested only on laser-targeted renditions and was significant compared to laser shuffled data (Extended Data Fig. 4a,b). Suppression of LMAN-RA terminals also reduced variability measured across all acoustic features (Extended Data Fig. 4c), consistent with prior experiments^28,29^.

As with the sBG suppression, the regressive effect of either HVC-sBG or LMAN-RA terminal suppression on syllable maturity was consistent throughout the day (Extended Data Fig. 4d) and suppressing either HVC-sBG or LMAN-RA terminal activity tended to most strongly affect individual acoustic features that were aligned with syllable maturity (Fig. 2f,g). Suppressing HVC or LMAN terminals in the sBG, or LMAN terminals in RA, did not alter variability of syllable maturity, syllable syntax, inter-syllable gap duration, or affect the ultimate similarity of syllables shared by the tutor and pupil (Extended Data Fig. 4e-h). However, unlike sBG cell body suppression, the regressive effects of HVC-sBG or LMAN-RA terminal suppression on syllable maturity remained constant across sensorimotor learning (Extended Data Fig. 4i). Collectively, these experiments identify HVC-sBG and LMAN-RA synapses as sites where juvenile sensorimotor learning is initially expressed.

### A role for cortico-sBG synapses in imitative learning

A hallmark of sensorimotor learning is its imitative nature: juveniles learn in reference to the song of an adult tutor^4,5^. In contrast, the syllable maturity metric is entirely self-referential, as it quantifies learning by assessing syllable quality in relation to the individual’s ultimate performance without consideration of its similarity to the tutor. To determine whether the cortico-sBG circuitry also contributes to imitative aspects of sensorimotor learning, we retrained the VAE using both tutor and pupil songs, then calculated the distance in latent space between the pupil’s syllable renditions and the tutor syllable that served as its model (Extended Data Fig. 5a). This analysis revealed that many syllables became more similar to their tutor models across learning, as evidenced by a reduced distance in the latent space, consistent with an imitative process. However, other syllables failed to converge on— and in some cases, diverged from—the tutor model (Extended Data Fig. 5b,c). This lack of convergence could reflect precocious maturation prior to the start of our audio recordings, inaccurate tutor song memorization, poor sensorimotor learning, or competing processes that favor individuation, such as song innovation or improvisation^30,31^.

To assess the contributions of cortico-sBG circuitry to imitative learning, we focused our re-analyses of optogenetic experiments on those syllables that displayed an orderly convergence on the tutor song over the period of sensorimotor learning studied here (79/164 syllables, see methods). Suppressing either the sBG or HVC-sBG terminals—but not LMAN-sBG or LMAN-RA terminals—immediately and transiently increased the distance to tutor (Extended Data Fig. 5d-h). Transient optogenetic suppression of any of these terminals did not affect final learning outcomes, as assessed by the average distance between syllables shared by the tutor and pupil on the final recording day (Extended Data Fig. 5i). Collectively, these analyses show that activity at the HVC-sBG synapse is necessary for expressing rapid imitative changes during sensorimotor learning.

### Activity of both cortical inputs in the sBG is required for learning acquisition

The model tested here posits that coincident, singing-related activity of HVC and LMAN terminals generates a synaptic eligibility trace in the sBG that is subsequently reinforced by a dopamine (DA) signal that evaluates syllable quality^15^. Consistent with this model, in the juvenile finch, DA dynamics in the sBG track syllable maturity^26,32^, and blocking D1-type DA receptors in the sBG prevents syllable maturation^26^ and tutor song copying^33^. However, while the optogenetic experiments performed here indicate that the sBG and specifically HVC-sBG synapses are necessary for the expression of recent learning, they are insufficient to establish the site of learning acquisition. For example, activity at LMAN-RA terminals is also necessary to express recent sensorimotor learning (Fig. 2e), raising the possibility that learning is a distributed process or occurs downstream of the sBG.

To test whether learning acquisition requires activity at both HVC-sBG and LMAN-sBG terminals but not at LMAN-RA terminals, we expressed inhibitory G_i/o_-coupled hM4Di chemogenetic receptors^34^ (H4) bilaterally in either HVC or LMAN of juvenile male finches and implanted infusion cannulae bilaterally in the sBG or RA (Fig. 3a; HVC-sBG n = 4 birds, LMAN-sBG n = 4 birds, LMAN-RA n = 5 birds). We recorded song production from days 60 to days 70 and focally infused the H4-agonist Clozapine-n-Oxide (CNO, 1.0 mM) into the sBG or RA on day 67 (Fig. 3b). At the end of this recording period, we anesthetized a subset of these birds and made electrophysiological recordings to confirm that systemic CNO treatment reduced spontaneous sBG or RA action potential activity for several hours only in H4-expressing birds (Fig. 3a, Extended Data Fig. 6a,b). Additionally, in another set of juvenile birds (n = 3), we co-expressed axon-targeted GcAMP6s and hM4Di receptors in LMAN neurons and confirmed that infusion of CNO focally to RA strongly attenuated singing-evoked Ca^2+^ transients in LMAN-RA terminals (Extended Data Fig. 6c,d).

**Figure 3.**
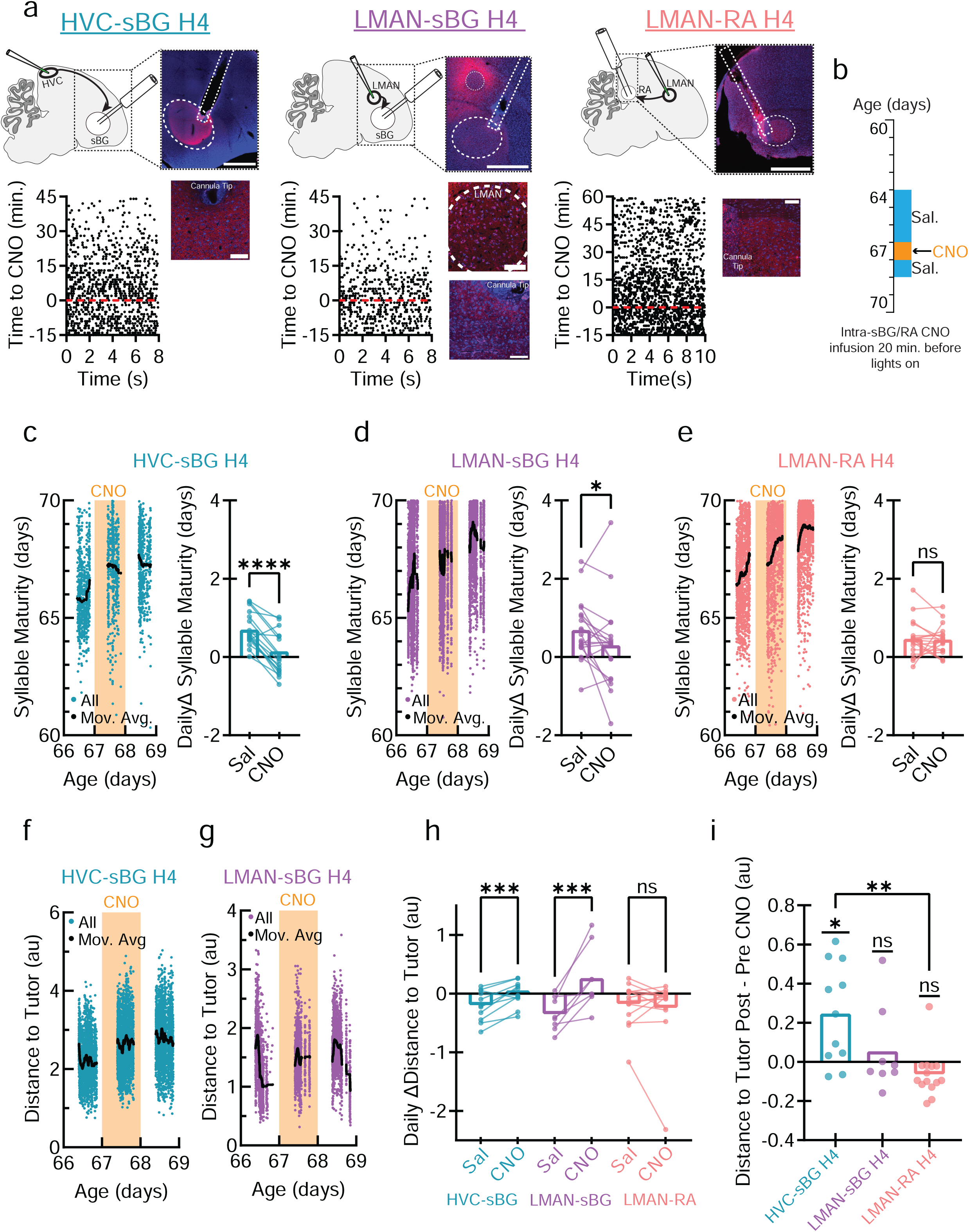
Either HVC-sBG or LMAN-sBG—but not LMAN-RA—terminal activity is necessary for learning acquisition. (**a**) (top) Schematics and representative histology (scale bar = 1 mm) from terminal chemogenetic suppression experiments (red = H4-mCherry, blue = Neurotrace, dashed lines = cannula and region of interest). (bottom) Higher magnification images of sBG/RA terminal fields (scale bars = 100 µm) and example raster plots (50% subsampled) showing reduced sBG/RA activity after systemic H4-agonist Clozapine-n-oxide (CNO) injection. (**b**) Experimental timeline. (**c**) (left) Syllable maturity (Mov. Avg. = 200 rendition Moving Average) for example HVC-sBG H4 syllable. (Right) Linear mixed effects model predicting daily change (Δ) in Maturity during Saline (Sal) or CNO treatment found a significant Treatment x Daily Change interaction (F_1,164994_= 29.2, p = 6.66e-8). (**d**) As in (**c**) but for the LMAN-sBG H4 group (F_1,165241_ = 4.33, p = 0.037). (**e**) As in (**c**) but for the LMAN-RA H4 group, no CNO effect detected (non-significant interaction: F_1,145387_ = 0.02, p = 0.88). (**f**) Distance to Tutor for HVC-sBG H4 example syllable. (**g**) As in (f) for an LMAN-sBG H4 example. (**h**) Linear mixed effects models predicting daily change in Distance to Tutor found a significant Treatment x Daily Change interaction in the HVC-sBG H4 group (F_1,98817_ = 11.7, p = 0.0006) and LMAN-sBG H4 group (F_1,62043_= 18.2, p = 1.99e-5), but not the LMAN-RA H4 group (F_1,146249_ = 0.39, p = 0.52). (**i**) Change in Distance to Tutor from the day before CNO treatment to the final recording day. The HVC-sBG H4 group *increased* in distance to the tutor (one-sample t-test t_10_ = 3.22, corrected p = 0.027). A Kruskal-Wallis test found a significant group difference (H = 12.1, p = 0.002). Dunn’s corrected post hoc tests revealed a difference only between HVC-sBG and LMAN-RA (Z = 3.47, p = 0.002). H4 = inhibitory hM4Di chemogenetic receptor. au = arbitrary units. * = p < 0.05, ** = p < 0.01, *** = p < 0.001, **** = p < 0.0001. ns = not significant.

At the end of the recording period (after day 70), we constructed VAE-ANN models to quantify syllable maturity (∼1.3 million syllable renditions). Suppressing activity with focal CNO treatment at either HVC-sBG or LMAN-sBG terminals—but not LMAN-RA terminals—abolished or strongly blunted daily increases in syllable maturity (Fig. 3c-e). Furthermore, for those syllables that approached the tutor over the course of sensorimotor learning (33/58), focally suppressing HVC-sBG or LMAN-sBG terminals—but not LMAN-RA terminals—either prevented or reversed the normal daily progress towards the tutor syllable (Fig. 3f-h). In addition, suppressing HVC-sBG terminals with focal CNO treatment for a single day led to a persistent increase in the separation between pupil and tutor syllables on subsequent days, an effect that was not observed in the LMAN-sBG or LMAN-RA groups (Fig. 3i). In contrast, daily increases in syllable maturity were unimpeded by CNO treatment in H4-expressing juveniles where the cannulae were inadvertently placed outside the sBG (n = 2 each for HVC-sBG and LMAN-sBG groups), showing that focal CNO treatment near but outside the sBG does not diffuse to HVC-sBG terminals, LMAN-sBG terminals, or LMAN cell bodies, or exert systemic effects that interfere with syllable maturation (Extended Data. Fig. 6e).

Lastly, while no manipulation altered syllable syntax, chemogenetically suppressing LMAN-sBG terminals reduced the variance of syllable maturity, and suppressing either LMAN-sBG or LMAN-RA terminals—but not HVC-sBG terminals—also modestly reduced variability measured across all acoustic features (Extended Data Fig. 6f-h).

These experiments establish that learning acquisition requires the activity of both HVC and LMAN terminals in the sBG, while activity at LMAN-RA terminals is necessary to express but not to acquire this recent learning.

### Augmenting sBG spiny neuron activity can both facilitate and disrupt song copying

Notably, the average maturity of syllables in fluorophore-expressing control birds in the ArchT experiments increased during laser illumination (Fig. 1j, Ctl: +1.99 ± 0.56 hours (mean ± SEM)), and laser administration could increase sBG action potential activity, likely because of tissue heating^35^ (Extended Data Fig. 7a). These results prompted us to test the possibility that artificially augmenting action potential activity in sBG spiny neurons could accelerate changes in syllable maturity. We expressed either the excitatory G_g_-coupled chemogenetic hM3Dq^34^ receptor (H3, n = 7 birds total; n = 5 for systemic CNO injection; n = 2 for intra-sBG CNO infusion via cannula), or fluorophore control (GFP, n = 6) in sBG spiny neurons of juvenile male finches using a CaMKIIa promoter^21^ (Fig. 4a; note that the CaMKIIa promoter also prevented H3 expression in LMAN; Extended Data Fig. 7b). We recorded song production from day 60 to day 70 and applied the H3-agonist CNO either systemically or focally in the sBG on day 67 during this period of sensorimotor learning (Fig. 4b). At the end of the recording period, we verified that systemic injection of CNO (1.0 mg/kg, i.m.) significantly increased sBG action potential activity for several hours in H3-expressing but not GFP-expressing animals (Fig. 4a, right; Extended Data Fig. 7c).

**Figure 4.**
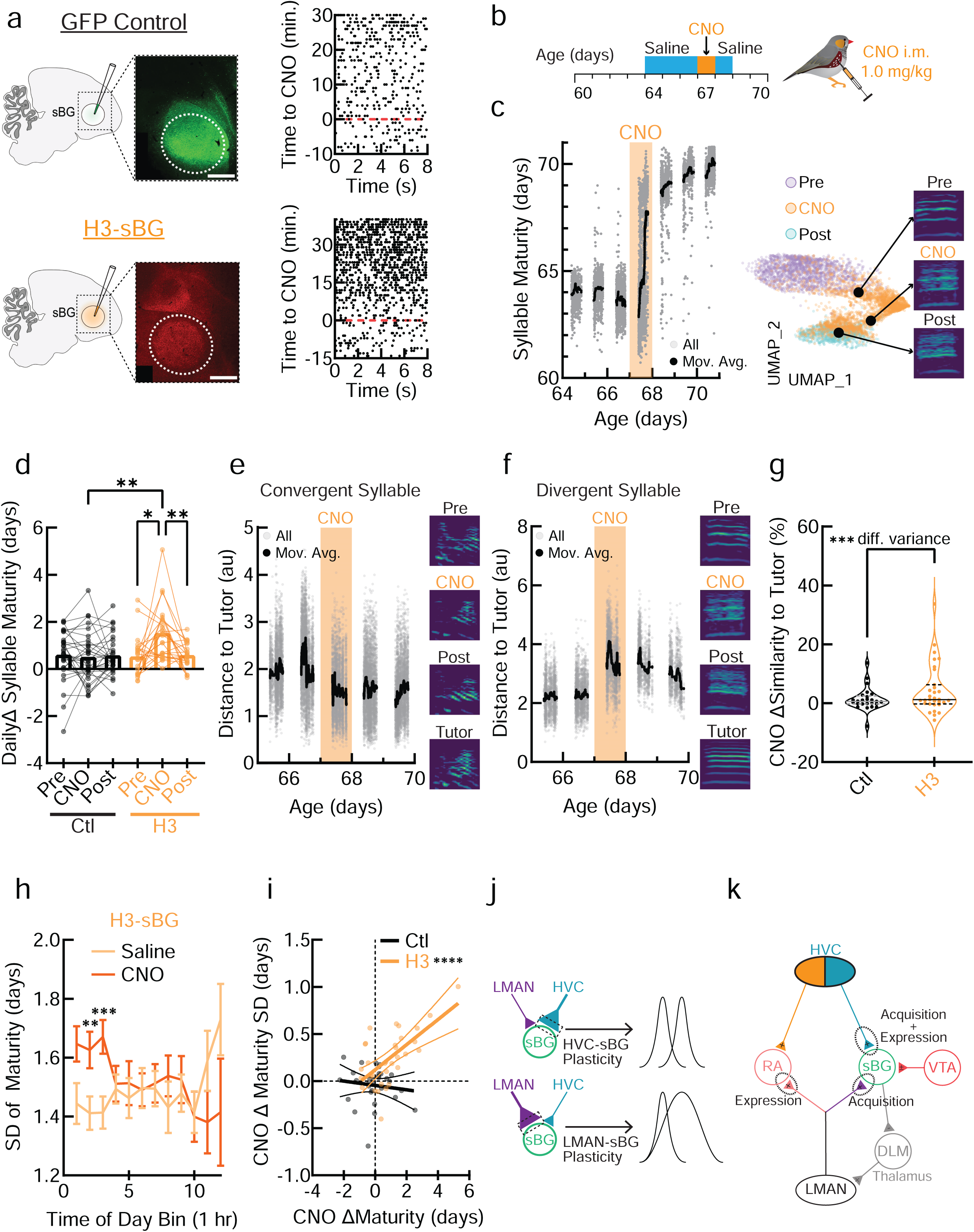
Transient activation of sBG spiny neurons accelerates vocal changes. (**a**) (Left) Schematic and representative histology for GFP Control (green) and H3-mCherry (red) expression in sBG spiny neurons. (Right) Example sBG raster plots (50% subsampled) relative to systemic injection of the H3-agonist Clozapine-n-oxide (CNO). (**b**) Experimental timeline (i.m. = intramuscular). (**c**) (Left) Syllable Maturity of all renditions and 200 rendition moving average (Mov. Avg.) in example H3 syllable, CNO day shaded in orange. (Right) UMAP projection and representative spectrograms from the days Pre, during, and Post CNO. (**d**) Daily change (Δ) in Maturity Pre, during, and Post CNO. 2-way RM ANOVA with Geisser-Greenhouse correction (Group x Treatment) found a main effect of Treatment (F_1.82,92.9_ = 5.16, p = 0.0093), and an interaction (F_2,102_ = 7.16, p = 0.0012). Corrected post hoc comparisons found that H3-CNO differed from pre- (t_23_ = 3.04, p = 0.0182) and post-treatment (t_23_ = 3.53, p = 0.0056), and Ctl-CNO (t_44.88_ = 3.17, p = 0.0027). (**e**) (Left) Distance to tutor (au = arbitrary units) for example H3 syllable that converged on the tutor and (right) representative spectrograms. (**f**) as in (e), only for an H3 syllable that diverged from the tutor. (**g**) Change in acoustic similarity to tutor from the day before CNO to 3 days post CNO. No difference in means, but different variance (F_24,30_ = 4.64, p = 0.0003). Solid line = median, dotted line = quartiles. (**h**) Standard Deviation (SD) of Maturity binned across the day for the H3 group during Saline/CNO. Mixed effects model with Geisser-Greenhouse correction revealed only a Treatment x Time of Day interaction (F_2.78, 63.7_ = 3.58, p = 0.021). Corrected post hocs identified significant differences during hours 2 (t_31_ = 4.02, p = 0.0041) and 3 (t_31_ = 4.64, p = 0.0007). (**i**) Linear relationship between CNO-induced changes in maturity and maturity SD (H3: F_1,30_ = 21.3, p < 0.0001, R^2^ = 0.415). (**j**) HVC-sBG plasticity would shift just the mean of maturity (post- relative to pre-plasticity), while LMAN-sBG plasticity would shift the mean and increase variance, which would be detrimental for expert performance. (**k**) The model tested here, which localizes the acquisition and expression of song learning to the HVC-sBG synapse, +/- indicate excitatory/inhibitory connections. Data points represent individual syllables unless noted otherwise. * = p < 0.05, ** = p < 0.01, *** = p < 0.001, **** = p < 0.0001. ns = not significant.

In juveniles expressing the H3 receptor, syllable maturity rapidly increased when sBG spiny neuron activity was augmented by CNO using either a single systemic injection (Fig. 4c,d) or intra-sBG infusion (Extended Data Fig. 7d-f; these groups were collapsed in subsequent analyses). Across syllables, these rapid increases in syllable maturity were not attributable to changes in syllable syntax (Extended Data Fig. 7g) or to changes of any single acoustic feature, suggesting that augmenting spiny neuron activity affected different acoustic dimensions in different syllables (Extended Data Fig. 7h). Thus, artificially elevating sBG spiny neuron activity is sufficient to accelerate syllable maturation.

For individual syllables, CNO-induced changes could be either convergent (Fig. 4e) or divergent (Fig. 4f) with respect to the pupil’s tutor model. Consequently, at the group level, CNO treatment did not change the average acoustic similarity to the tutor (Fig. 4g). Furthermore, even when restricting our analysis to tutor-convergent syllables, CNO treatment did not change the average distance to tutor in latent space (Extended Data Fig. 7i). However, CNO treatment did expand the range of learning outcomes, as reflected in a greater variability in both the acoustic similarity and latent distance to tutor metrics (Fig. 4g; Extended Data Fig. 7i; i.e., H3-expressing juveniles exhibited both larger *increases* and larger *decreases* in acoustic similarity and latent distance to tutor). Moreover, despite the transient nature of the CNO application, these maturational changes persisted for days and were readily apparent in representative spectrograms of syllables generated before and after treatment (Fig. 4c,e,f). Therefore, chemogenetically augmenting spiny neuron activity persistently altered the juvenile’s syllables without consistently driving them closer to their tutor model.

An influential concept is that sensorimotor learning relies on vocal exploration followed by selective reinforcement^22,29,36^. In this scenario, artificially augmenting spiny neuron activity would accelerate syllable maturation by elevating vocal variability, leading to more extensive vocal exploration. In fact, CNO treatment in H3-expressing juveniles increased the rendition-to-rendition variability of both syllable maturity (Fig. 4h) and acoustic features (Extended Data Fig. 7j). Moreover, this increased variability in syllable maturity positively correlated with the magnitude of subsequent changes in syllable maturity (Fig. 4i), as expected of an exploration-generating reinforcement learning mechanism.

## Discussion

Here we identified the specific synapses that enable the learning of birdsong, a complex, culturally-transmitted motor skill with key parallels to human speech and music^1^. We established that activity at the HVC-sBG synapse is necessary for both the acquisition and expression of recent sensorimotor learning in the juvenile finch.

Furthermore, we distinguished this dual role of the HVC-sBG synapse from other sites important only for learning acquisition (LMAN-sBG terminals) or expression (LMAN-RA terminals). We capitalized on these insights to further show that transient augmentation of postsynaptic sBG spiny neuron activity is sufficient to accelerate syllable maturation and induce enduring changes in song, and to determine that the sBG regulates vocal exploration in a flexible manner that aligns with both daily and longer-term learning goals.

Along with prior studies showing that DA dynamics in the sBG encode syllable quality and drive syllable maturation^26,32^, the current observations support a model^15^ in which coincident, singing-related activity of HVC and LMAN inputs to the sBG generates a synaptic eligibility trace in the sBG, which DA acts on to reinforce HVC-sBG synapses and thus drive sensorimotor learning^33,37,38^. Further, whereas activity at LMAN-sBG terminals is necessary to acquire but not express sensorimotor learning, activity at LMAN-RA terminals is necessary for learning expression but not acquisition, a double dissociation that is particularly striking given the same LMAN cell bodies give rise to both projections^11^. This parallels and extends studies in adult birds that distinguish between a role for the sBG in the acquisition of externally-reinforced pitch learning^33,37^, and a more restricted role for LMAN-RA synapses in its expression^28^.

While HVC axons form synapses on other, less abundant sBG cell types^39^ that may afford additional sites of plasticity, suppressing HVC axon terminals and sBG spiny neurons regressed maturity to a similar degree, identifying the HVC-spiny neuron synapse as a crucial locus for the expression of sensorimotor learning. This synapse’s critical role in learning is underscored by retrospective electron microscopy analyses of HVC-sBG spiny neuron synapses, which identify structural correlates of plasticity^40^, and the finding here that transient chemogenetic inhibition of HVC-sBG terminals, but not LMAN-sBG or LMAN-RA terminals, caused long-lasting deficits in song copying (Fig. 3i). These persistent deficits are noteworthy given that juvenile zebra finches ultimately display normal tutor song copying even after D1 DA receptors in the sBG are blocked for ten straight days during the middle of sensorimotor learning^33^. Restricting plasticity to HVC-sBG synapses presumably affords a temporally precise premotor “bias” signal while avoiding increased song variability that could arise from potentiating LMAN-sBG synapses (Fig. 4j).

Beyond identifying synaptic components necessary to acquire and express recently learned changes to the juvenile’s song (Fig. 4k), the current study sheds light on the timescale and logic of this complex learning process. Notably, suppressing either HVC-sBG terminals, sBG spiny neurons, or LMAN-RA terminals caused song maturity to regress several hours, and these effects were remarkably constant across the day and unaffected by overnight sleep. An inference is that plasticity within the sBG contributes to relatively short-lasting changes in song that are continuously consolidated at a relatively fast timescale (hours) elsewhere in the brain, likely at HVC-RA synapses^16,17^. The sBG’s contribution to song also waned with development, pointing to a slower developmental process wherein progressive consolidation ultimately renders song performance independent of the sBG^7,9,10^. Finally, optogenetically suppressing sBG activity modulated syllable features depending on the magnitude and sign of their contributions to daily changes in syllable maturity. Apparently, the sBG flexibly biases song performance rather than rigidly specifying movement parameters.

Although birdsong learning is sometimes characterized as rote copying^41^, an intriguing observation here is that sensorimotor learning in zebra finches could divergence from the tutor model, reminiscent of vocal improvisation and innovation described in other songbirds^42^. While chemogenetically elevating sBG activity increased song variability and rates of syllable maturation, these effects failed to enhance the overall quality of tutor song imitation. Whether this chemogenetic augmentation of syllable maturation involves the same exploration and reinforcement mechanisms of natural learning remains to be determined. However, one possibility is that activity levels in the sBG are normally titrated to optimize imitation, such that artificially augmenting sBG activity triggers persistent synaptic and vocal changes without improving overall imitation.

Another possibility is that imitation is only one of several goals of a learning process that also rewards individuation^30,31^ and that chemogenetic elevation of sBG activity enables the juvenile to explore and consolidate vocal variants otherwise outside of its normal capacity.

## Materials and Methods

### Animals

74 juvenile male zebra finches (*Taeniopygia guttata*) and their respective tutors (37 adult males) from the Mooney Lab breeding colony at Duke University Medical Center were used across experiments. Juveniles were kept on a 14:10 light:dark cycle and had *ad libitum* access to food and water. Prior to sound recording, juveniles were kept in their home cage with their parents and any siblings. All experiments were performed under a protocol approved by Duke University Institutional Animal Care and Use Committee.

### Surgical Procedures

Juvenile male zebra finches were anesthetized with 2% isoflurane, head-fixed on a stereotaxic device, and placed on a heating pad. Three successive rounds of alcohol/betadine, followed by 0.25% bupivacaine as a local anesthetic were applied to the scalp prior to making an incision. All experiments, except for calcium imaging experiments, were done bilaterally. Viruses were obtained from Addgene and the PPO plasmid from Addgene was transformed into a vector by the Duke Viral Vector Core.

Craniotomies were made to target regions relative to the Y-sinus as follows: sBG (coordinates chosen to avoid LMAN): head angle 43° 5.1 mm anterior, point marked, head moved to 72°, 1.2-1.5 mm anterior of mark, 1.6 mm lateral, -3.1, -2.8, -2.6 mm ventral (200 - 300 nl of virus at each depth for optogenetic experiments. Chemogenetic excitation experiments used a single injection of 150 nl at -2.9). LMAN: head angle 50°, 4.9 mm anterior, 1.5 mm lateral, -2.0, -1.8 mm ventral (100 nl each depth). HVC: head angle 18°, 0.1 mm anterior and 0.1 mm posterior, 2.35 mm and 2.45 mm lateral, -0.4 mm ventral (250 nl each site). Viruses were injected via a Nanoject-II (Drummond Scientific) in 10 nl pulses separated by 15 s. The pipette was left in place for 5 min (deepest injection) or 10 min (shallowest injection site) to allow for diffusion.

Craniotomies were closed with the bird’s own skull tissue, and the surgical site was sealed using Vetbond. Birds received Meloxicam as an analgesic and were monitored in a recovery cage with food, water, and a heat lamp until fully recovered before being returned to the bird colony or a sound attenuating chamber.

### Optogenetic Suppression

For the sBG suppression experiments, juvenile male zebra finches (20 - 35 days post hatch or dph) were injected with a virus expressing a control fluorophore (AAV2/1-CAG-tdTomato, n = 4), or the inhibitory opsin Archearhodopsin (ArchT) (AAV2/1-CaMKIIa-ArchT-GFP (n = 3) for spiny neuron specific expression^21^ and AAV2/1-CAG-ArchT-GFP (n = 3) for pan-neuronal expression) targeting the sBG. After allowing for 3 weeks of viral expression (41 - 56 dph), birds were implanted with bilateral tapered optical fibers (3 mm total length with 1.0 mm taper, Optogenix) targeting the sBG and secured using Kwik-Sil (WPI) and Metabond (Parkell, S396, S398, S371). Prior to implantation, an optrode was lowered into the sBG, and we pulsed 2s of constant 532 nm laser (GL532T3-300FC, Shanghai Lasers) while recording neural activity and laser timestamps (Differential A-C Amplifier 1700, A-M Systems). We compared sBG activity during the 2s of laser illumination to activity in the preceding 2s. Birds without apparent optogenetic suppression of sBG firing were not implanted with optical fibers (an additional n = 3 ArchT birds).

For the terminal suppression experiments, juvenile male zebra finches (20 - 35 dph) were injected with a virus expressing the G_i/o_-coupled opsin Parapinopsin^27^ (PPO, AAV2/9-CAG-PPO-mVenus) targeting HVC (n = 5) or LMAN (n = 5). PPO was used to avoid paradoxical excitation seen with terminal expression of many traditional ion channel opsins^43^. Five fluorophore control birds were injected with a GFP-expressing virus (AAV2/9-CAG-eGFP) targeting HVC. As above, we implanted tapered optical fibers bilaterally in the sBG (i.e., the terminal field) after 3 weeks (41 - 56 dph) and verified suppression in the sBG using an optrode. We pulsed 462 nm laser at 20 Hz with 5-10 ms pulses for 2 - 5s (BLM462TA-200FC, Shanghai Lasers). Note that while recent reports suggest PPO responds primarily to shorter, near UV wavelengths, 462 nm lasers were not filtered and produced equivalent power at the shorter wavelengths necessary to activate PPO in these new reports^43^. In an additional n = 2 PPO birds, we failed to identify responsive units and so these birds were not implanted with fibers.

In 2 HVC-sBG birds (one experimental bird and one used just for this purpose), we stimulated HVC with a bipolar stimulating electrode (WE5ST30.1H10, MicroProbes) using a single 1 ms, 500 µA pulse. In alternating blocks of 10 trials, we then either only administered the electrical stimulation, or we preceded and followed the electrical pulse with 1s of laser. We performed electrical stimulation experiments only in the HVC group because a substantial fraction of HVC axons pass through LMAN on their way to the sBG^13,44^ such that LMAN electrical stimulation would activate both of the sBG’s cortical inputs, and because the proximity of LMAN to the sBG (<1.0 *mm*) compared to HVC (∼1.0 *cm*) renders simultaneous stimulation in LMAN and recording in the sBG technically challenging.

For LMAN-RA terminal suppression, juvenile male zebra finches (25 - 35 dph) were injected with the same PPO (n = 4) or GFP (n = 4) viruses in LMAN, and bilateral tapered optical fibers were implanted in RA (coordinates: 35°, 1.35 caudal, 2.45 lateral, -2.5 V) after allowing at least 2 weeks for expression. Prior to implantation, we recorded RA activity and verified PPO-mediated suppression of RA firing using an optrode.

For all optogenetic groups, after recovery from the bilateral fiber implantation surgery, birds were individually housed in sound isolating boxes (∼60 dph), tethered to optical fibers, and we began recording all the vocalizations they produced over the course of a month (∼90 dph) using Evtaf software^22^. After allowing the number of songs to stabilize over the course of 2-5 days, we implemented our transient suppression of motifs. We designed a template for each bird to target the introductory notes of his song motif, and then, three days each week (MWF) we used Evtaf software^22^ to detect introductory notes and trigger 2 s of laser (constant 532 nm for ArchT; 20 Hz 5-10 ms pulses, 462 nm for PPO) on a random 10% of motifs with less than 10 ms of jitter. The laser could not turn on again for at least 3 s following stimulation. Experimental and control birds received laser at similar powers (∼10 mW) to control for non-specific effects of heating and/or light.

### Chemogenetic Suppression

For the chemogenetic suppression experiments, juvenile male zebra finches (30 - 45 dph) were injected with a virus expressing the inhibitory G_i/o_-coupled hM4Di designer receptor exclusively activated by a designer drug (DREADD)^34^ in either HVC or LMAN (AAV2/9-hSyn-hM4Di-mCherry, n = 6 HVC and n = 11 LMAN). In the case of the HVC injections, all birds were implanted with bilateral drug infusion cannulae targeting HVC terminals in the sBG (5.0 mm long, 3.0 mm separation, Protech International). In the case of the LMAN injections, n = 6 birds were implanted with bilateral cannulae targeting LMAN terminals in the sBG, while n = 5 birds were implanted with bilateral cannulae targeting LMAN terminals in RA (two, single, 5.0 mm long cannula, Protech International). Birds were given 2 weeks prior to the start of behavioral recordings and at least 3 weeks before we administered the selective H4-agonist Clozapine-n-Oxide (CNO, water soluble variant, Thermo SML2304). Following the completion of behavioral procedures, we recorded sBG or RA action potential activity in anesthetized birds and verified that CNO (1.0 mg/kg 20 µl in saline, intramuscular or i.m.) reduced sBG or RA firing (n = 3 LMAN-RA birds, n = 2 LMAN-sBG birds, and n = 1 HVC-sBG bird), and had no effect in one additional control bird in RA or in the sBG (see subsequent chemogenetic excitation experiment, n = 1 bird).

After surgical recovery, birds were individually housed in sound isolating chambers from ∼60 dph until 70 dph, and all vocalizations were recorded using Sound Analysis Pro software^45^ (http://soundanalysispro.com). Dummy cannula (Protech, 0.5 mm projection from guide cannula) were left in place until we began intra-sBG/RA infusions, when we switched them out for internal cannula (Protech, 0.5 mm projection). Vehicle (Ringer’s solution) was infused from dph 64-66 to habituate birds to handling and control for non-specific effects of infusion. CNO was infused on dph 67 (1.0 mM in Ringer’s), followed by a final vehicle day on dph 68. We infused 250 nl of fluid (sBG) or 500 nl (RA) at a rate of 0.5 µl/min (Harvard Apparatus, 70-3007) and left internal cannula in place for 1 minute for diffusion. Infusions were completed 20 minutes prior to lights on to ensure that birds recovered and to give enough time for CNO to act prior to the start of singing (∼10-15-minute latency based on our recordings of sBG/RA activity).

In n = 2 birds each from the HVC-sBG and the LMAN-sBG experiments, cannulae were inadvertently placed outside of the sBG and were excluded from the main analyses.

However, their data was used as a control for non-specific effects of CNO and to assess CNO spread. That is, although there was H4 expression in HVC/LMAN terminals in the sBG in these birds, cannula placement ∼800 µm outside the sBG was insufficient to elicit a behavioral effect (Extended Data Fig. 6e), suggesting CNO does not diffuse to LMAN cell bodies (slightly greater ∼1000 µm distance) when the cannula was properly targeted to the sBG.

### Optofluid LMAN-RA Calcium Imaging

We used viral vectors to express both an axon-targeted GCaMP6s (AAV2/9-hSyn-axon-GCaMP6s-P2A-mRuby3) and an inhibitory chemogenetic receptor hM4Di (AAV2/9-hSyn-hM4Di-mCherry) in LMAN (200 nl at -1.9 and -1.8 mm ventral, 1:1 ratio of viruses). We then implanted an optofluid cannula (4.0 mm fiber with 200 µm core, 3.8 mm guide cannula, Doric Lenses) unilaterally above LMAN’s terminal field in RA (n = 5 juvenile birds total with n = 2/5 birds excluded due to optofluid cannulae placement outside of RA) to enable us to both measure LMAN-RA terminal calcium transients and focally deliver the H4-agonist CNO to RA. After waiting 2 weeks for viral expression, birds were tethered to a patch cord (Doric Lenses, 200 µm core, 0.37 NA) and imaged using a photometry system (Neurophotometrics FP3002). Briefly, (see also^26^) light from a 470 nm LED to excite GCaMP6s fluorescence was alternated with a 415 nm LED isosbestic control at 15 Hz per wavelength. LEDs were bandpass filtered, passed down the patch cord via a 20X objective, and light output for each wavelength was set to ∼50 µW at the tip of the optical fiber. Emitted GCaMP6s calcium fluorescence was collected through the same patch cord, split by a dichroic mirror, bandpass filtered and focused onto a sCMOS camera. Photometry and vocal data were acquired using Bonsai software^46^.

Photometry data were analyzed using custom MATLAB (Mathworks) scripts. In brief, the blue channel was fit to and subtracted from the green channel using MATLAB’s *polyfit* function. Song bout onsets were labeled, and the green calcium signal was Z-scored relative to a baseline period (-4 to -2 s) prior to song onset. After several 5-minute-long baseline recording sessions were acquired, birds were disconnected from the patch cord, dummy cannula were removed, and 500 nl of Vehicle (Ringer’s solution) or CNO (1.0 mM) was infused into RA at a rate of 500 nl/minute. The injection cannula was left in place for 1 minute after infusion, after which the dummy was reinserted, the bird was re-attached to the patch cord, and data was periodically recorded in ∼5-minute-long sessions.

### Chemogenetic Excitation

For the chemogenetic excitation experiments, juvenile male zebra finches (30 - 45 dph) were injected with a virus expressing the excitatory G_q_-coupled hM3Dq DREADD^34^ under the CaMKIIa promoter to target sBG spiny neurons (AAV2/9-CaMKIIa-hM3Dq-mCherry, n = 5) or a fluorophore control (AAV2/9-CaMKIIa-eGFP, n = 6). Birds were given 2 weeks prior to the start of behavioral recordings, and at least 3 weeks before we administered the CNO. Following the completion of behavioral procedures, we recorded sBG action potential activity in anesthetized birds and verified that CNO (1.0 mg/kg 20 µl in saline, intramuscular or i.m.) increased sBG firing in two H3-expressing birds and had no effect in one control bird. Behavioral procedures and timelines matched those used in the chemogenetic suppression experiments, with the exception that birds received intramuscular (i.m.) injections of saline (20 µl) and CNO (20 µl, 1.0 mg/kg) rather than intracranial infusions.

In two additional birds, we injected the H3-expressing virus into the sBG and implanted bilateral cannulae targeting the sBG in the same surgery. Experimental procedures mirrored those of the chemogenetic suppression experiments, although we injected a larger amount of fluid (1 µl) since we were not concerned with CNO spread to LMAN cell bodies since the virus was injected into the sBG rather than LMAN, and since our use of a CaMKIIa promoter precluded expression in LMAN (Extended Data Fig. 7b).

### Tissue Collection and Immunofluorescence

Following the completion of behavioral procedures, birds were deeply anesthetized with intramuscular injections of Euthasol (20 µl, Virbac) and transcardially perfused first with 1X phosphate-buffered saline (PBS, Sigma P5493) then 4% paraformaldehyde (PFA). Brains were removed, post-fixed in 4% PFA overnight at 4°C, then moved to 30% sucrose PFA as a cryoprotectant for two days. Brains were sectioned on a cryostat (Leica CM 1850) at 50 - 60 µm, collected into 1X PBS, and stored at 4°C. For immunofluorescence, we collected 4 - 6 sections per hemisphere targeting our region of interest. Free-floating sections were first washed three times for five minutes in PBST (0.3% Triton X-100, Sigma, T8787), then blocked with 10% Blocking One Histo (Nacalai tesque, 06349-64) in PBST for 1 hour. Sections were incubated overnight with primary antibody, washed 3 times for five minutes each wash in PBST and then incubated in secondary antibody with Neurotrace 435/455 (1:500, Invitrogen, N21479) for four hours at room temperature. Sections were mounted and coverslipped with Fluoromount-G (SouthernBiotech, 0100-01) and imaged under a confocal microscope (Zeiss 710).

To localize expression of ArchT-GFP, PPO-mVenus, axon-GCaMP6s and eGFP, we used mouse anti-GFP primary (1:500 Invitrogen, A11120). To localize the expression of tdTomato, we used rabbit anti-RFP primary (1:1000, Rockland, 600-401-379). To localize the expression of H3- or H4-mCherry, we used rabbit anti-mCherry primary (1:500, Abcam, AB167453). To localize expression of CaMKIIa protein, we used rabbit anti-CaMKIIa primary (1:500, Abcam, AB22609). For all mouse primaries, we used goat anti-mouse Alexa Fluor 488 secondary (1:500, Invitrogen, A11001). For all rabbit primaries, we used goat anti-rabbit Alexa Fluor 594 secondary (1:500, Invitrogen, A11032). We assessed viral expression in all birds and excluded any animals with insufficient or off-target expression. For all optogenetic and cannula experiments, we also ensured that fibers/cannulae targeted the sBG/RA and were proximate to viral expression.

### Variational Autoencoder (VAE)

After recording a juvenile’s vocalizations across most of sensorimotor learning (∼60-90 dph using Evtaf (32 kHz) for optogenetic experiments, ∼60-70 dph using Sound Analysis Pro (44.1 kHz) for chemogenetic experiments), we segmented songs into their component syllables using an amplitude-based segmentation pipeline in the Autoencoded Vocal Analysis python package^18^. These segments were used to generate spectrograms via a short time Fourier Transform (time segments: 512 samples, overlap: 256), with the amplitude appropriately scaled and the frequency range clipped for each bird to exclude noise. These segment-level spectrograms were interpolated to 128 x 128 linearly spaced frequencies and time points. Spectrograms were then fed into a standard Variational Autoencoder (VAE) model, as described previously^18,19^. Briefly, the VAE was trained separately for each bird using 50000 spectrogram segments for 100-200 epochs when loss stopped decreasing on the validation set. The latent dimensions were fixed at 32, and all parameters were optimized using Adam optimization with a learning rate of 0.001%. We used Uniform Manifold Approximation and Projection (UMAP) to generate a reduced representation of the 32 VAE latent embeddings for visualization and used a custom MATLAB (Mathworks) script to plot the 2 UMAP dimensions and assign syllable labels to well-clustered groups. We plotted at least 40 randomly-selected examples for each putative syllable cluster and excluded syllables with an error rate greater than 5% (i.e., different syllables and or noise included within the putative syllable cluster).

### Artificial Neural Network (ANN)

For each syllable, we trained an Artificial Neural Network (ANN) with 3 linear layers to predict the age at which each syllable rendition was produced based on the 32 latent embeddings, as described previously^19^. Syllable ages were scaled between -1 to 1 for each syllable, and network parameters were optimized by minimizing the sum of squared errors between a syllable’s predicted age of production and the actual age of production using the Levenberg-Marquardt algorithm and Bayesian regularization. We used 80% of a syllable’s renditions as a training set with the remaining 20% set aside for model validation and stopped training if there was no improvement in the validation set for 3 epochs, or until 30 minutes had elapsed. Importantly, in the optogenetic experiments, all laser-targeted renditions were excluded from ANN training. For the chemogenetic experiments, both the preceding vehicle treatment day, and the CNO treatment day were excluded from training.

### Syllable Maturity Calculations

To assess effects of optogenetic suppression, we built linear mixed effects models, described in more detail below. Syllable maturity was centered each day by subtracting the daily mean of syllable maturity from all renditions on a given day. To assess potential circadian effects, we calculated a locally-centered version of syllable maturity by aligning to each laser-targeted rendition and calculating the maturity difference between that rendition and the average of the nearest hour (30 minutes before/after) of non-laser targeted renditions. We then calculated the average of this local syllable maturity difference within daily 3 hour-long bins. To assess potential developmental changes in laser efficacy, we calculated the average laser-induced change in day-centered syllable maturity each day, then fit a linear regression relating these laser effects to age. To determine if the effects of optogenetic suppression were transient, we calculated the average difference in day-centered syllable maturity for the current rendition (n) when the laser occurred *only* on a specific preceding rendition or “lag” (e.g. only on n - 3) for each syllable type. For the excitatory chemogenetic experiments, we calculated the change in maturity from the start to the end of the day (last 50 renditions - first 50 renditions). For the inhibitory chemogenetic experiments, we used linear mixed effects models (described below) to model the daily slope of syllable maturity. Standard deviation (SD) of maturity was calculated within 1 hour-long bins, and we performed a linear regression to relate changes in maturity SD (CNO - Vehicle) to daily changes in syllable maturity (CNO - Vehicle) in Fig. 4i.

### Distance to Tutor Analysis

The distance to tutor VAE was trained similarly to the pupil-only VAE, except that now both pupil and tutor songs (recorded in the same sound-attenuating chamber) were included in the training. We again used UMAP to assign syllable labels to clusters, ensuring that these clusters included both the tutor and the pupil. Consistent segmentation of syllable types was more challenging in these joint models, leading to different numbers of identified syllables in the Distance to Tutor datasets. Some syllables in the pupil did not have clear analogues in the tutor song or could not be consistently classified and were therefore not included in this analysis. We then calculated the mean latent representation of each tutor syllable (a single 32-dimensional point) and calculated the Euclidean distance between this tutor mean and every rendition of that same syllable made by the pupil. We built syllable-level linear models to predict distance to tutor as a function of days post hatch (dph), time of day, and laser. A negative dph coefficient was used to classify a syllable as “convergent” with the tutor (i.e., the syllable linearly approached the tutor across learning). Syllables with non-significant or significantly positive dph coefficients were excluded as “divergent” syllables. In the excitatory chemogenetic experiment, we made this inclusion decision based on the days after CNO treatment (dph 68-70), since CNO treatment could sometimes shift a syllable from convergent to divergent (Fig. 4f) or vice versa (Fig. 4e). We excluded divergent syllables throughout all experiments because the putative goal of song learning - the tutor model - is no longer clear. Distance to tutor is a metric of learning only when we can be reasonably certain that the tutor model was the pupil’s goal; in “divergent” syllables, distance to tutor is orthogonal to actual learning.

### Linear Mixed Effects Models

To determine if Laser influenced Syllable Maturity against a background of age-related changes in maturity, we built linear mixed effects models (*fitlme* function in MATLAB) where we modeled Syllable Maturity as follows:

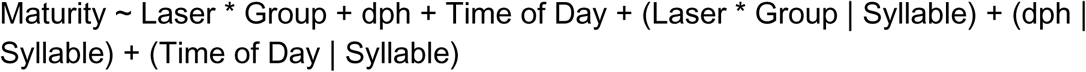

Where * indicates an interaction and main effects, Laser is a categorical variable indicating if Laser did or did not target a rendition, and parenthetical terms indicate modeling of random effects at the Syllable level. The Laser * Group interaction and the effects of Laser within each group were assessed with linear hypothesis tests to determine if there were significant Group differences and/or significant effects of Laser within each Group. Tests were post hoc corrected, and degrees of freedom were determined using the residual method, but significance was unchanged when using the Satterthwaite Approximation.

To determine if Laser effects differed from order shuffled data, we shuffled the Laser variable within syllable and within day (since Laser only occurred on certain days) and trained linear mixed effects models on data from 1000 Laser shuffled datasets. These models were trained individually for each group (i.e., one model for Control and another for Experimental) due to the extensive training time required for the previously described combined Group Interaction models that precluded performing the necessary number of permutations. Significant effects were assessed using post hoc corrected, two-tailed permutation tests of the form:

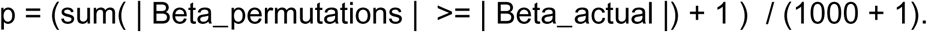

In the chemogenetic suppression experiments, we built linear mixed effects models as follows:

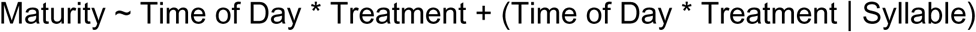

Where * indicates an interaction and main effects, Treatment is a categorical variable comparing the CNO treatment day vs the preceding three saline/vehicle treatment days, and parenthetical terms indicate modeling of random effects at the Syllable level. We specifically tested the Time of Day * Treatment interaction term to see if CNO treatment significantly affected the daily slope of maturity relative to the preceding saline days.

### Elastic Net Regressions

To determine if and how acoustic features related to maturity, and if and how laser affected individual acoustic features we built elastic net regression models (MATLAB’s *lasso* function) using data only from days with laser administration (3 days/week). We built two versions of these models, one trained at the syllable level (one model/syllable) and another trained at the day level (one model/syllable/day). Syllable maturity and the predictors (acoustic features) were z-scored separately for each syllable, or within each syllable, each day respectively for these two model types. These elastic net regressions minimize mean-squared error while penalizing the sum of absolute values of the parameters using a regularization weight, which we set, *a priori*, to 0.5. This penalizes predictor variables that do not provide a unique contribution to predicting the outcome variable and pushes their coefficients to zero, an effect which is useful when predictors are multicollinear, as is the case for acoustic features. We used 5-fold cross validation to select LASSO lambda values such that the mean squared error was within one standard error of the minimum. We repeated the elastic net fitting 10 times for each model and took the average of the coefficient values to ensure stable coefficients.

The Maturity as a function of Feature models (Maturity ∼ Feature) yielded coefficients for each of the acoustic features. The Feature ∼ Laser models predicted one feature at a time, with all other features included as covariates (e.g., Mean Frequency ∼ Laser + all other features), and we report the coefficients relating Laser to a specific feature. To examine the relationship between Laser-effected Feature changes and how those Features related to Maturity, we took the vector of Feature coefficients for the Maturity ∼ Feature models and compared this to the vector of Laser coefficients for the Feature ∼ Laser models using cosine similarity. That is, we asked whether the collection of Laser effects on Features aligned with the collection of Features that linearly predicted increases in Maturity. We also repeated this cosine similarity calculation within the model types across syllables or across days to assess the stability of the sets of Features that contribute to Maturity, and the stability of Laser effects on Features across syllables and days.

### Song Feature Analysis

Using the syllable labels generated by the VAE, we segmented raw audio files into individual syllables. We then used Sound Analysis Pro’s “Batch” analysis method to quantify a set of 13 standard acoustic features^45^ for each rendition of each syllable, separately for syllables that were or were not targeted by laser (optogenetic experiments, using only days where laser was administered) or on days with Saline vs. CNO treatment (chemogenetic experiments). These data were exported into Excel (Microsoft) spreadsheets and imported into MATLAB for analysis. We calculated the coefficient of variation (CV; SD / Mean) of acoustic features and compared this CV value on Laser renditions relative to non-laser renditions (Laser On / Laser Off * 100%). Since Laser occurred on a small subset of renditions (∼10%) and to ensure that our CV measure was not biased by sample size, we resampled the Laser Off renditions to have the same sample size as the Laser renditions, calculated the Laser Off CV on each of 1000 permutations, and then took the median of the 1000 permutations to use as the Laser Off CV. For the chemogenetic experiments, we calculated the CV of each acoustic feature on the CNO day relative to the pre-treatment vehicle day (CNO / Vehicle * 100%). We compared Groups and Feature CV values using 2-way ANOVAs. In addition to the “Batch” analysis, we also used Sound Analysis Pro’s “*SAT_sound*” function in MATLAB to calculate a set of 15 standard acoustic features directly from our syllable segmented wav files. This dataset was used to align the acoustic feature data with the syllable maturity datasets to calculate relationships between features and maturity (see previous section on elastic net models). We also calculated pitch and mean frequency contours from the SAT_sound function, and normalized them as per^36^ to give us 17 features in total that we related to maturity and to effects of laser.

To complement our VAE-based Distance to Tutor measurement, we also calculated a more classic measure of similarity to tutor that is based on a syllable’s acoustic features^45^. To calculate acoustic similarity to tutor, we again used the VAE labels to segment both pupil and tutor song into their respective syllables. We used Sound Analysis Pro’s Similarity Batch analysis to calculate the acoustic similarity to tutor as a percentage for all the pupil renditions from the final recording day (optogenetic experiments) to a tutor syllable exemplar. For the chemogenetic excitation experiment, we calculated the Acoustic Similarity score one day before and 3 days after treatment with CNO and calculated the change in Acoustic Similarity. We note that Distance to Tutor is a more robust measure than Acoustic Similarity since it uses the average tutor rendition rather than an exemplar and quantifies all pupil renditions. Indeed, the Similarity Batch could rarely (4/217 syllables) fail to quantify similarity for syllables that Distance to Tutor identified as convergent.

### Syllable Syntax

Using the segmentation decisions of the VAE, we calculated the syllable transition matrix. We found the proportion of “canonical” transitions for each syllable (i.e., A>B vs. A>any other syllable). We then calculated the proportion of canonical transitions for laser on versus off and compared this change in transition probability to 0 using a one-sample t-test, or an unpaired t-test/one-way ANOVA to assess group differences.

## Statistical Analysis

All analyses were two-tailed with α = 0.05 as the threshold for significance. Post hoc tests were corrected for multiple comparisons using the Bonferroni method. Data were analyzed using MATLAB (Mathworks) and Prism (Graphpad).

## Data Availability

The datasets generated in the current study will be uploaded to Duke University Library Research Data Repository pending instructions from the Editor. They are currently available from the corresponding author on request.

## Code Availability

The code used for the variational autoencoder and artificial neural network are available at: https://github.com/pearsonlab/autoencoded-vocal-analysis and https://github.com/SamuelBrudner/juvenile_syllable_analysis, respectively.

## Acknowledgements

The authors thank Steve Lisberger, Christina Gremel, Miles Martinez, Jiaxuan Qi, and Kate Kaplan for comments on an earlier draft of this manuscript. This research was supported by NIH F32 MH132152 and NIH K99 NS144525 to DS, NIH F31 HD098772 to SB, NIH 5R01 NS099288 to RM and RF1 NS118424 to RM and JP.

## Author contributions

Conceptualization: DCS, SB, RM, JP. Methodology: DCS, SB. Investigation: DCS, AL. Visualization: DCS. Formal Analysis: DCS. Funding acquisition: DCS, SB, RM, JP. Project administration: RM and JP. Supervision: RM and JP. Writing – original draft: DCS and RM. Writing – review & editing: DCS, SB, AL, RM, JP.

## Competing interests

Authors declare that they have no competing interests.

## Supplementary Information

No additional supplementary information.

## Correspondence

Correspondence and requests for materials should be addressed to RM.

---

**Extended Data Figure 1.**
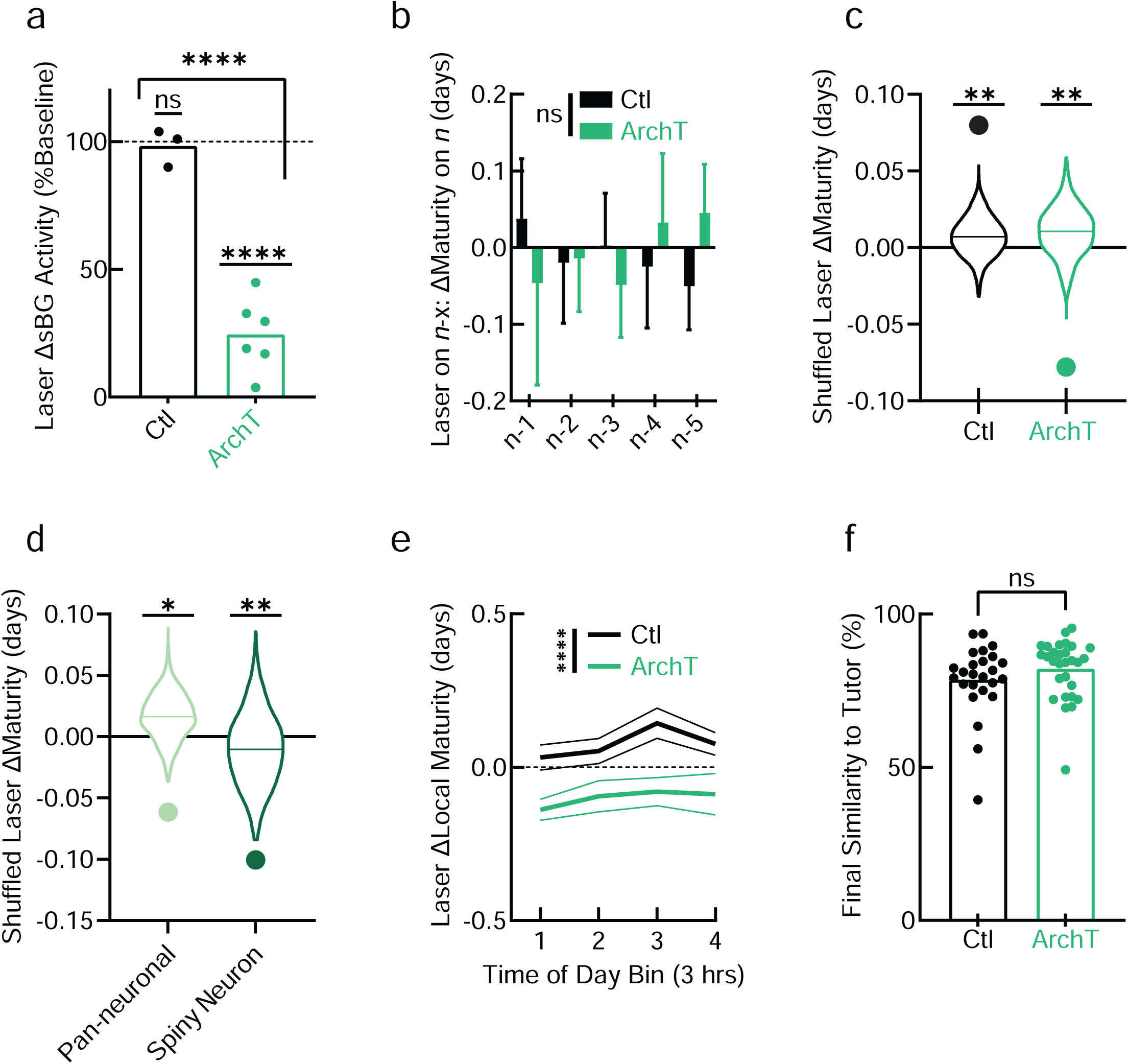
sBG suppression reliably reduces maturity. (**a**) Average (over 10 trials) laser-induced change (Δ) in sBG activity (% of 2s pre laser). sBG activity was reduced only in ArchT (unpaired t-test: t_7_ = 8.20, p < 0.0001; one-sample t-test vs. 100%: t_5_ = 12.9, p < 0.0001). Points represent one unit from one bird (3/4 Ctls recorded). (**b**) Change in syllable maturity of a given syllable type during rendition *n* when laser occurred only on the preceding *n* - x rendition, no significant effects detected (2-way RM ANOVA). (**c**) Laser coefficients from 1000 Laser shuffled mixed effects models (violin plots, solid line = median) and actual Laser coefficients (points). Post hoc corrected permutation tests found significant differences from shuffled data in Ctl (p = 0.002) and ArchT (p = 0.002). (**d**) As in (c) but showing that Laser effects were significant relative to shuffled data in both pan-neuronal (p = 0.02) and spiny neuron-specific (p = 0.002) ArchT expression groups. (**e**) Local maturity (see methods) grouped in 3-hour bins, 2-way RM ANOVA detected a main effect of group only (F_1, 47_ = 22.8, p < 0.0001) and not time of day nor an interaction. (**f**) Acoustic similarity to tutor on the final recording day, points = syllables. Bars are Mean ± SEM. * = p < 0.05, ** = p < 0.01, **** = p < 0.0001. ns = not significant.

**Extended Data Figure 2.**
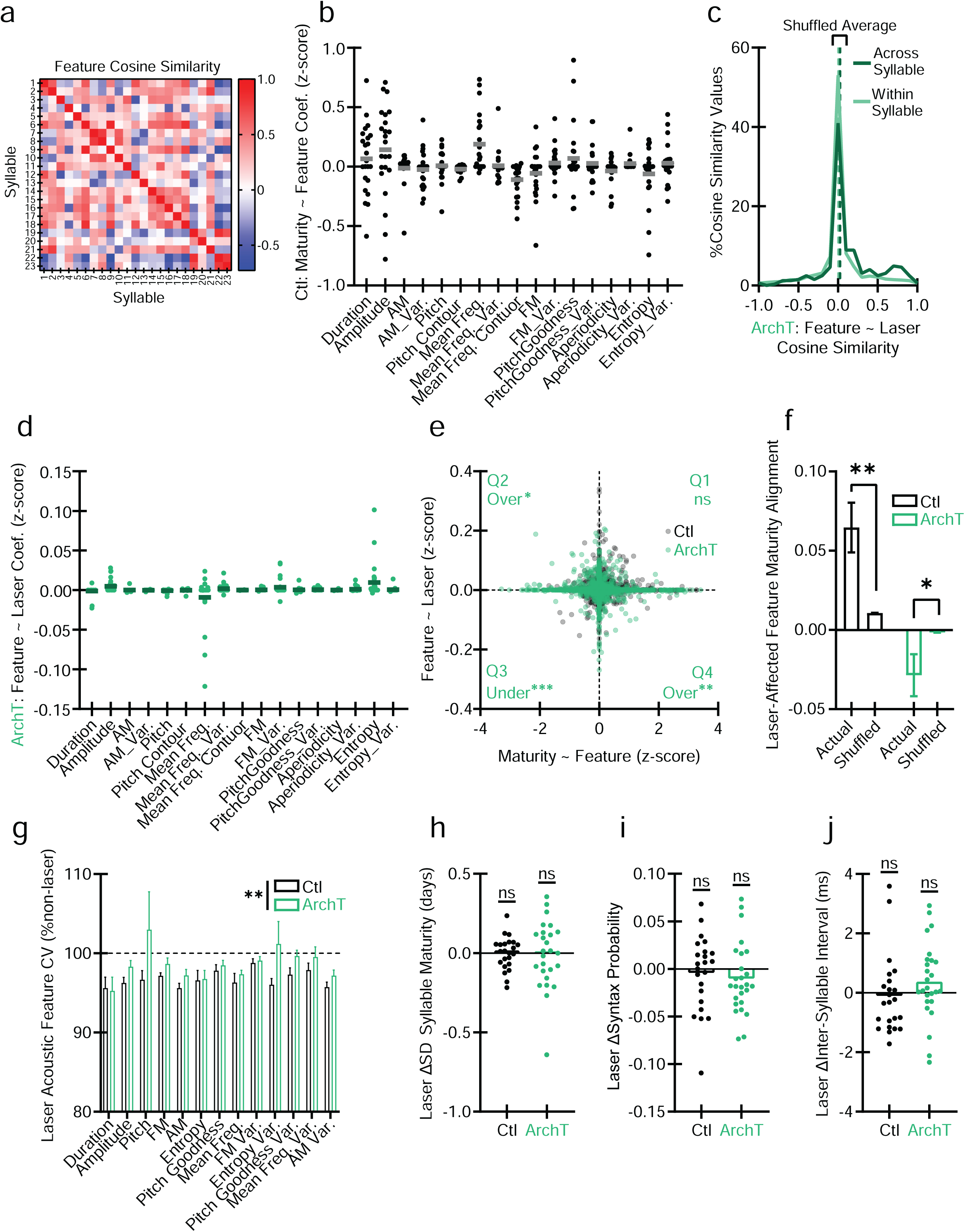
sBG suppression tends to affect acoustic features when they align with maturity. (**a**) Cosine similarity matrix comparing Feature vectors from elastic net regressions predicting Maturity as a function of Acoustic Features (Maturity ∼ Features) across 23 Control syllables. (**b**) Maturity ∼ Feature coefficients (Coef.) for individual acoustic features in the Ctl group. Points represent syllables, bars = across syllable mean. Substantial variability in the presence, direction, and magnitude of these relationships. (**c**) Histogram showing the distribution of Feature ∼ Laser cosine similarities in the ArchT group Across Syllables or Within Syllables but across days. (**d**) Feature ∼ Laser coefficients for individual acoustic features in ArchT. Laser makes small, inconsistent contributions to individual features. (**e**) Scatter plot relating Maturity ∼ Feature vs. Feature ∼ Laser coefficients in Ctl and ArchT shows that quadrants 2 and 4 are over-represented in ArchT (quadrant 2: Z = 2.25, p = 0.025; quadrant 4: Z = 2.8634, p = 0.0042; quadrant 3 under-represented: Z = -3.42, p = 0.00062), indicating that, when a feature predicts maturity (horizontal axis), laser tends to push that feature in the opposite direction (vertical axis). Points represent the relationship of one syllable and one feature on a single day (e.g., Syllable A, Mean Frequency, day 75). (**f**) Alignment of laser-affected features with features that predict maturity. Cosine similarity of actual and shuffled data between the vector of acoustic features that predict maturity and the vector of laser effects on those same features. Positive numbers indicate that the effect of laser is aligned with maturity in feature space, while negative numbers indicate anti-alignment. Actual/Shuffled difference in Ctl (KS-test: D = 0.097 n_1_ = 279, n_2_ = 279 x 10000 shuffles, p = 0.009) and ArchT (KS-test: D = 0.075 n_1_ = 345, n_2_ = 345 x 10000 shuffles, p = 0.04). (**g**) Change in the coefficient of variation (CV) of all acoustic features when the laser is on relative to off, (2-way ANOVA (Group x Feature) detected a main effect only of Group F_1,610_ = 10.5, p = 0.0013). (**h**) Change in standard deviation (SD) of maturity when laser is one relative to when it is off. (**i**) Change (Δ) in transition probability for the canonical (e.g. A→B) syllable sequence for laser on vs. off. (**j**) Laser change in inter-syllable intervals. No effects detected h-j. AM = Amplitude Modulation. Freq. = Frequency. FM = Frequency Modulation. Var = Variance. * = p < 0.05, ** = p < 0.01. ns = not significant.

**Extended Data Figure 3.**
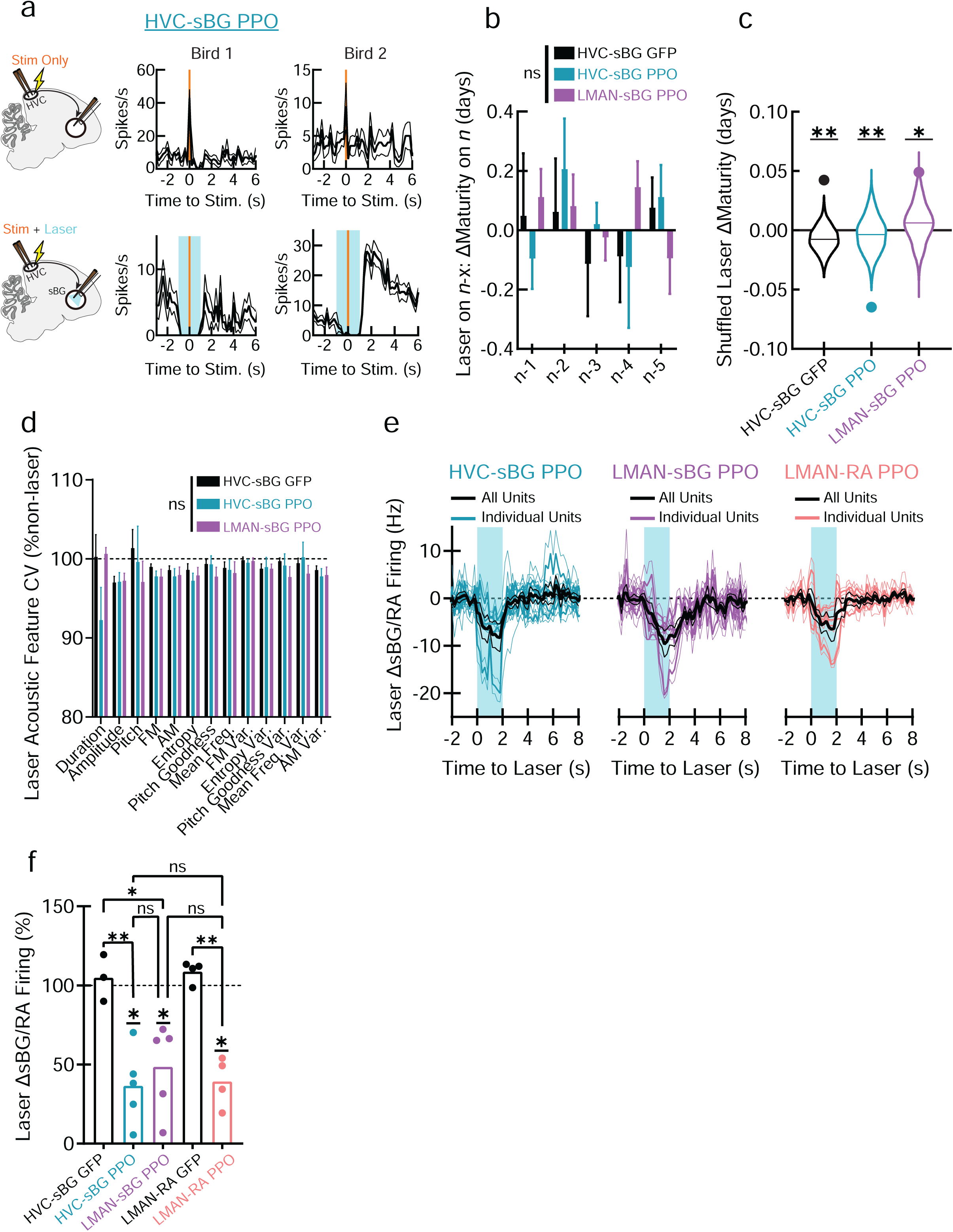
HVC-sBG suppression reliably reduces maturity. (**a**) (Top row) Electrical stimulation of HVC (1 ms pulse, 500 µamp, orange line) evokes postsynaptic action potential activity in the sBG. (Bottom row) Laser suppression (blue shaded region) of HVC-sBG PPO terminals prevents HVC-evoked activity in the sBG (1 unit/bird, 10 trials/condition). (**b**) Change (Δ) in syllable maturity of a given syllable type during rendition *n* when laser occurred *only* on the preceding *n* - x rendition, no significant effects detected (2-way RM ANOVA). (**c**) Mixed effects model shuffled Laser coefficients (violin plots, solid line = median) alongside the actual laser coefficients (points). There was a significant difference from shuffled data in each group, but maturity significantly regressed only in the HVC-sBG group (1000 shuffles, post hoc corrected permutation tests: HVC-sBG GFP p = 0.003; HVC-sBG PPO p = 0.003; LMAN-sBG PPO p = 0.015). (**d**) Change in the coefficient of variation (CV) of all acoustic features when the laser is on relative to off (2-way ANOVA (Group x Feature) detected no significant effects. (**e**) Change in sBG (left and center) and RA (right) activity (baseline mean subtracted) during laser-activation of PPO (blue shading) in cortical inputs. Individual Units averaged across 10 trials, All Units averaged across units. Thick/thin lines = mean/SEM. (**f**) Change in sBG/RA activity during 2 s of laser relative to preceding 2 s (one-way ANOVA: F_4,16_ = 12.1, p = 0.0001. Corrected post hoc comparisons: HVC-sBG GFP vs. LMAN-sBG PPO: t_16_ = 3.76, p = 0.01, HVC-sBG GFP vs. HVC-sBG PPO: t_16_ = 4.55 p = 0.002. LMAN-RA GFP vs. LMAN-RA PPO: t_16_ = 4.78, p = 0.0012. None of the PPO groups differed from one another). Corrected one-sample t-tests in PPO groups found significant effects in LMAN-sBG PPO (t_4_ = 4.09 p = 0.045), HVC-sBG PPO (t_4_ = 5.93 p = 0.012), and LMAN-RA PPO (t_3_ = 7.77 p = 0.013). Points = one unit from one bird (3/5 HVC-sBG GFP birds recorded). AM = Amplitude Modulation. Freq. = Frequency. FM = Frequency Modulation. Var = Variance. * = p < 0.05, ** = p < 0.01, ns = not significant.

**Extended Data Fig. 4.**
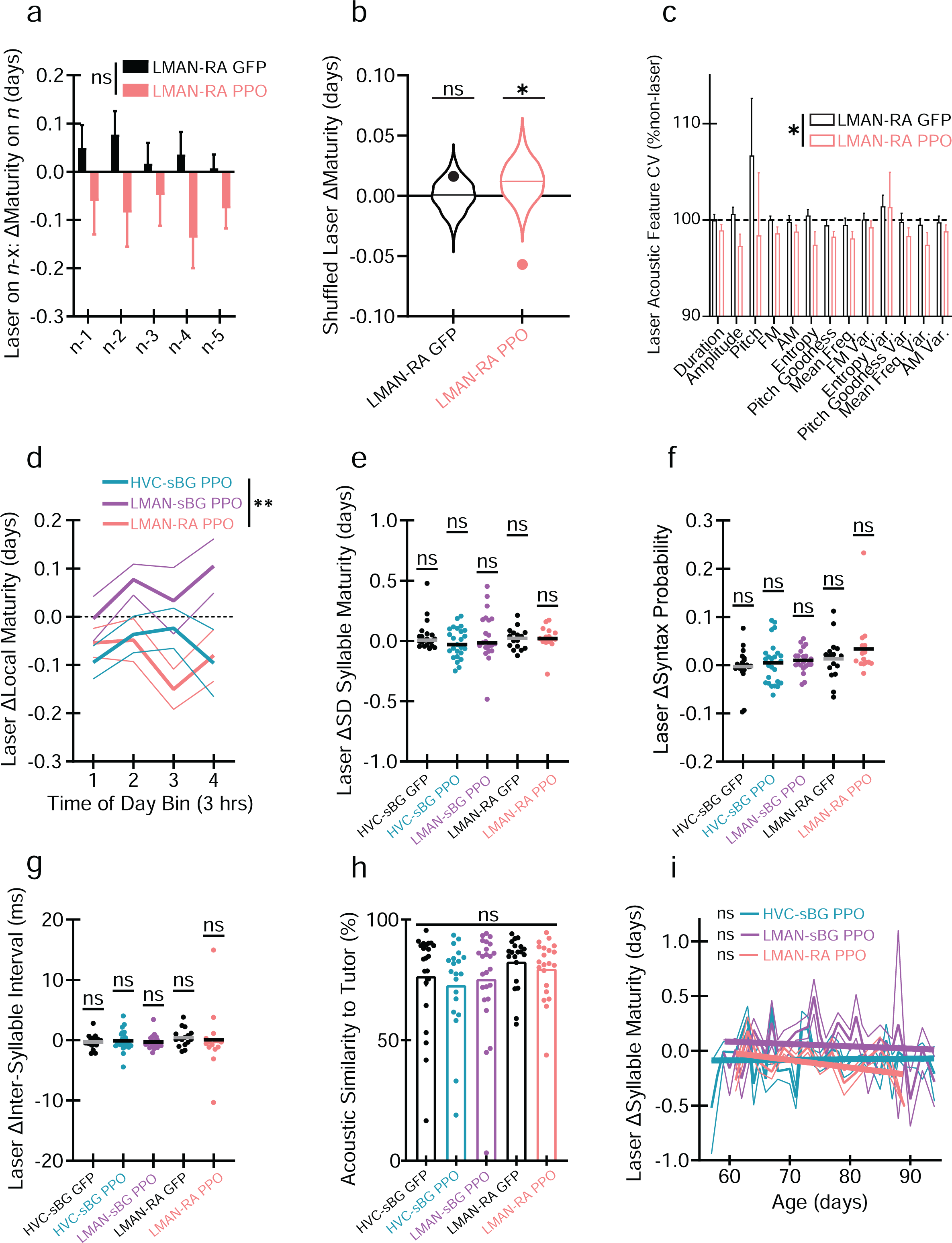
LMAN-RA suppression reduces syllable maturity and acoustic variability. (**a**) Change (Δ) in syllable maturity of a given syllable type during rendition *n* when laser occurred *only* on the preceding *n* - x rendition, no significant effects detected (2-way RM ANOVA). (**b**) Mixed effects model shuffled Laser coefficients (violin plots, solid line = median) and actual Laser coefficients (points). Only the LMAN-RA PPO group differed from shuffled data (1000 shuffles, corrected permutation p = 0.01). (**c**) LMAN-RA suppression reduced acoustic feature coefficient of variation (CV) (2-way ANOVA (Group x Feature) identified a significant effect only of Group F_1,377_ = 6.57, p = 0.011). (**d**) Change in Local Maturity (methods) grouped in 3-hour bins. 2-way (Group x Time of Day) RM Mixed Effects analysis revealed a main effect only of Group (F_2,58_ = 6.78, p = 0.002; corrected post hoc tests identified differences between LMAN-sBG PPO and both HVC-sBG PP (t_58_ = 3.12, p = 0.009) and LMAN-RA PPO (t_58_ = 3.19, p = 0.007). (**e**) Change in syllable maturity standard deviation (SD) when laser is on. (**f**) Change in transition probability for the canonical (e.g. A→B) syllable sequence for laser on vs. off. (**g**) Laser-induced change in inter-syllable interval. For (e)-(g) lines indicate mean values and neither one-way ANOVAs nor corrected one-sample t-tests identified significant effects. (**h**) No group differences in acoustic similarity to tutor on the final recording day (one-way ANOVA). (**i**) Average laser-induced change in Syllable Maturity grouped by day. Solid lines are a linear regression relating age to laser-induced changes in maturity. No slopes differed from 0. Points represent syllables unless otherwise specified. Bars = mean, lines in (a),(c),(d),(i) are SEM. AM = Amplitude Modulation. Freq. = Frequency. FM = Frequency Modulation. Var = Variance. * = p < 0.05, ** = p < 0.01, ns = not significant.

**Extended Data Figure 5.**
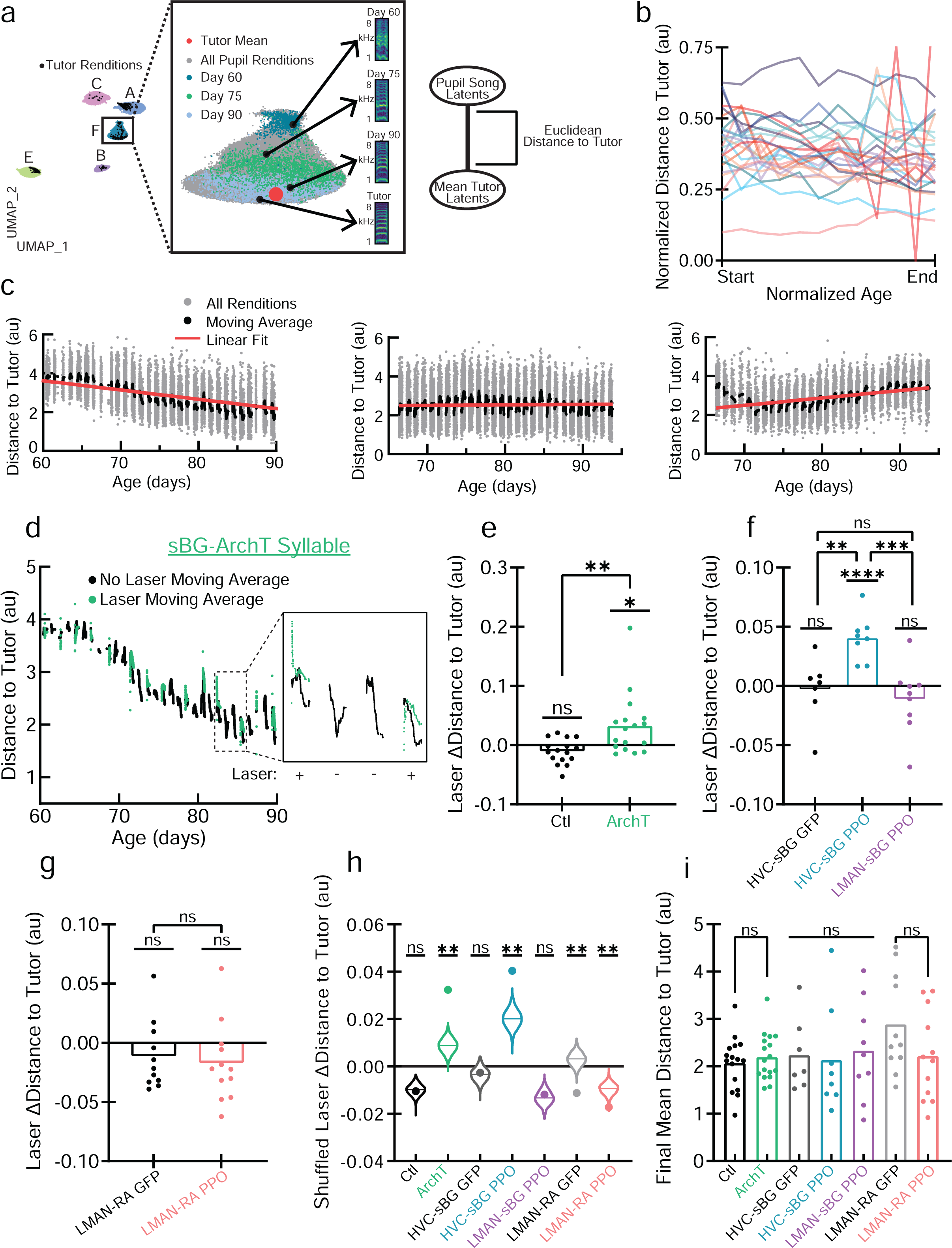
sBG and HVC-sBG activity is necessary to express imitative aspects of learning. (**a**) (left) A Variational Autoencoder (VAE) trained with both pupil and tutor songs places them into a shared latent space. UMAP representation of the VAE latent embeddings of pupil/tutor syllables reveals clusters. (center) Progression of syllable “F” from days 60-90 with all renditions colored in gray and specific days noted alongside representative spectrograms. The pupil renditions approach the tutor mean (red) in latent space across learning. (Right) Distance to tutor is the Euclidean distance in the VAE latent embeddings between each pupil rendition and the mean tutor rendition of that syllable. (**b**) Normalized distance to tutor and age (scaled between 0 to 1 and fit using spline approximations for visualization only) in the sBG ArchT group demonstrates the substantial syllable-to-syllable variability of change in distance to tutor during sensorimotor learning. (**c**) Example syllable patterns (200 rendition-long moving average) showing (left) orderly convergence on the tutor, (center) failure to approach the tutor, perhaps due to precocious maturation, and (right) initial approach followed by subsequent divergence from the tutor. (**d**) Distance to tutor with and without laser for an example sBG-ArchT syllable. Laser +/- indicates days with/without laser. (**e**) Change (Δ) in distance to tutor when laser is on estimated in a mixed effects model. Laser effects significantly differed between Ctl and ArchT (t_2.53e6_ = 2.94, p = 0.003), and post hoc corrected linear hypothesis tests identified significant effects only in ArchT (t_2.53e6_ = 2.49, p = 0.013). (**f**) as in (**e**) but for the HVC/LMAN-sBG suppression experiment. Corrected Laser effects significantly differed between HVC-sBG PPO and both HVC-sBG GFP (t_2.33e6_ = 3.23, p = 0.0036) and LMAN-sBG PPO (t_2.33e6_ = 3.90, p = 0.0003), and significant laser effects were identified only in HVC-sBG PPO (t_2.33e6_ = 4.95, p = 2.18e-6). (**g**) as in (**e**) but for LMAN-RA suppression. No significant laser effects (LMAN-RA GFP: p = 0.45, LMAN-RA PPO: p = 0.20) or group differences (p = 0.68) detected. (**h**) Shuffled mixed effects model Laser coefficients (violin plots, line = median) and actual Laser coefficients (points). Laser effects significantly differed from shuffled data (corrected permutation tests on 1000 shuffles) in ArchT (p = 0.002), HVC-sBG PPO (p = 0.003), LMAN-RA GFP (p = 0.008), and LMAN-RA PPO (p = 0.004). (**i**) Average distance to tutor on the final recording day. Groups compared within their respective experiments, no significant differences detected. Data points represent individual syllables unless otherwise specified. au = arbitrary units. * = p < 0.05, ** = p < 0.01, *** = p < 0.001, **** = p < 0.0001. ns = not significant.

**Extended Data Fig. 6.**
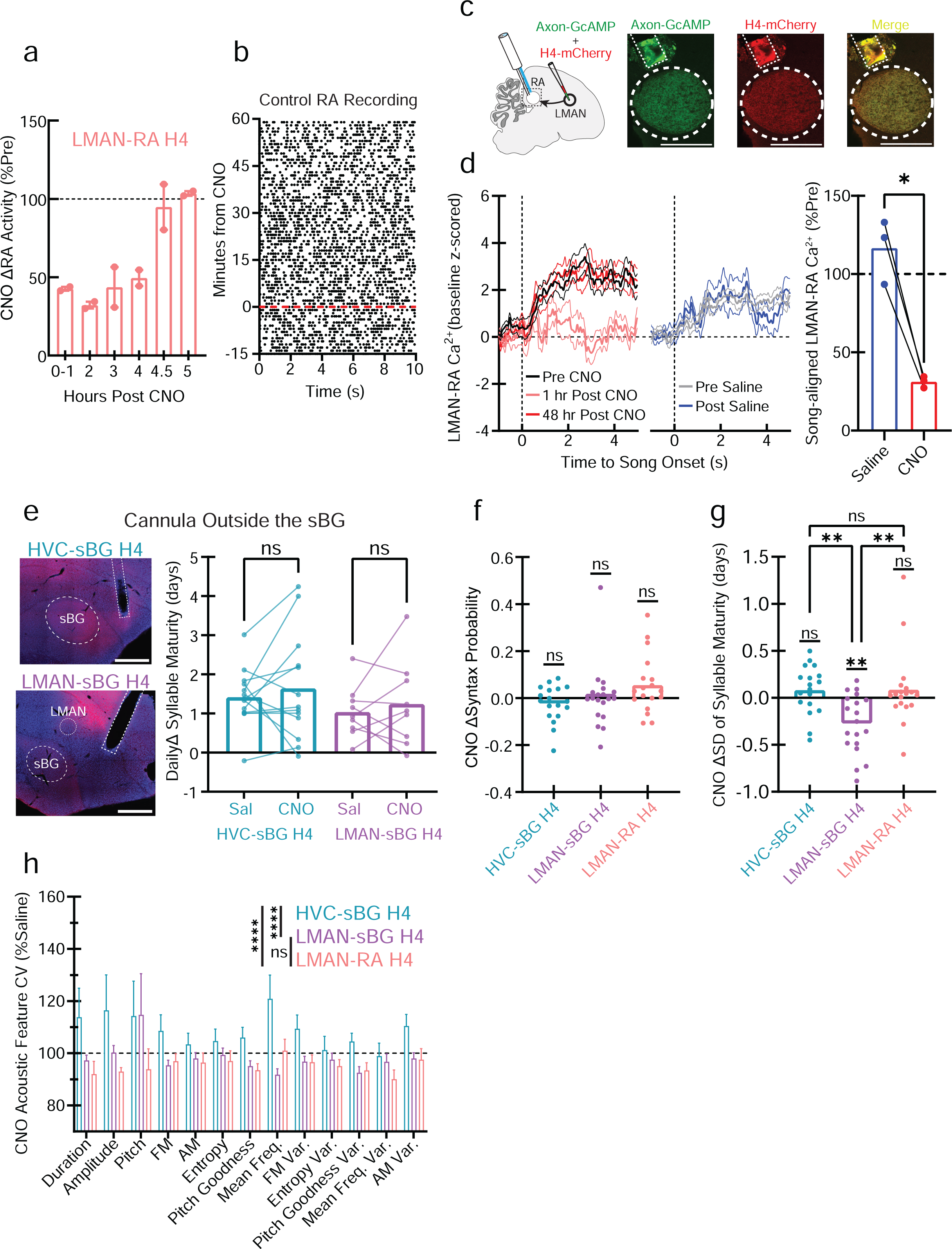
CNO effects on maturity are attributable to HVC-sBG and LMAN-sBG terminals and LMAN-sBG suppression affects maturity variability. (**a**) RA activity is reduced for ∼4 hours following systemic CNO injection, points = units. (**b**) Raster plot (50% subsampled) showing CNO does not affect RA activity in a control bird (no H4 expression). (**c**) Schematic and example histology showing optofluid cannula placement above RA and co-expression of Axon-GcAMP6s (green) and H4-mCherry (red) in LMAN-RA terminals (scale bar = 0.5 mm). (**d**) Average (+/- SEM) LMAN-RA terminal Ca^2+^ activity aligned to the onset of song bouts (averaged across bouts for one bird, z-scored relative to pre-song baseline) before and after focal CNO (left) or Saline (center) infusion. (right) Average LMAN-RA Ca^2+^ activity at song onset (0-2 s) following CNO or Saline (%pre-infusion baseline, points = birds). Significant Saline vs. CNO difference (paired t-test, t_2_ = 8.31, p = 0.013). (**e**) (left) Example histology (blue = Neurotrace, red = H4-mCherry, dashed lines = cannula) where cannula was placed outside the sBG (scale bar = 1.0 mm). (right) A mixed effects model predicting daily change (Δ) in maturity as a function of treatment found no significant CNO effects. (**f**) CNO-induced change (CNO - Saline) in the proportion of canonical syllable transitions (e.g., A→B), no effects detected. (**g**) CNO-induced change (CNO - Saline) in standard deviation (SD) of syllable maturity. Significant group difference (one-way ANOVA: F_2,52_ = 7.49, p = 0.0014), corrected post hoc tests identified differences between LMAN-sBG H4 and the other groups (HVC-sBG H4: t_52_ = 3.3, p = 0.005; LMAN-RA H4: t_52_ = 3.4, p = 0.004). Significant CNO effect only in LMAN-sBG H4 (corrected one-sample t-tests, t_18_ = 3.78, p = 0.004). (**h**) Coefficient of Variation (CV) of acoustic features on the CNO treatment day as %Saline. 2-way ANOVA (Group x Feature) identified a main effect of Group only (F_2,676_ = 20.1, p < 0.0001). Corrected post hoc tests identified a difference between HVC-sBG H4 and both LMAN-sBG H4 (t_676_ = 4,81, p < 0.0001) and LMAN-RA H4 (t_676_ = 5.99, p < 0.0001). Points = syllables in e-g. FM = Frequency Modulation, AM = Amplitude Modulation, Freq. = Frequency, Var = Variance. * = p < 0.05, ** = p < 0.01, **** = p < 0.0001. ns = not significant.

**Extended Data Figure 7.**
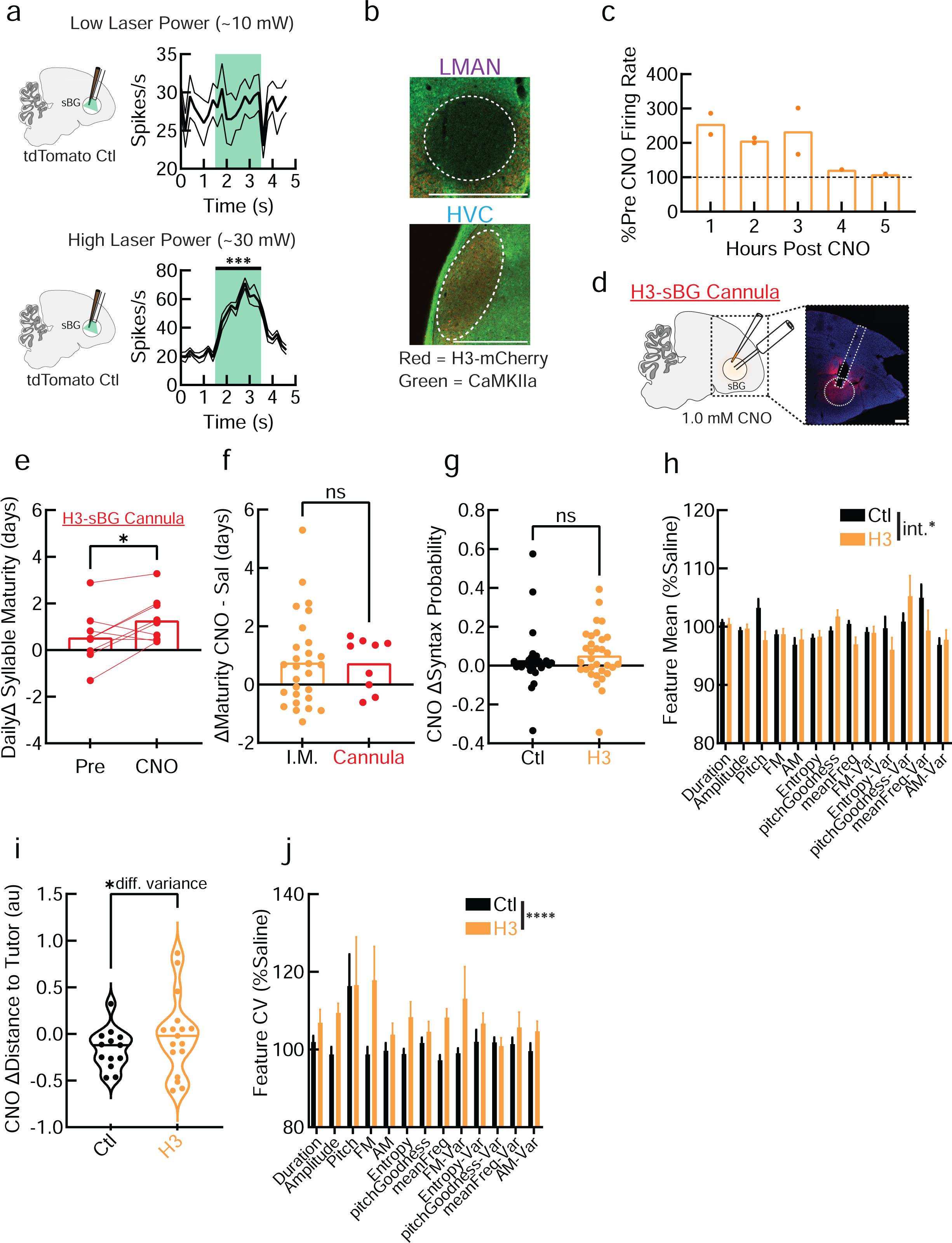
Intra-sBG CNO infusion accelerates song changes. (**a**) Example sBG recordings in one tdTomato control (Ctl) bird, green shading indicates constant 532 nm laser, showing increased activity with high power (paired t-test, t_7_ = 7.64, p = 0.0001). (**b**) Example histology in HVC and LMAN for a bird injected with AAV-CaMKIIa-H3-mCherry in the sBG, showing some retrograde infection (red) of HVC, but not LMAN due to lack of CaMKIIa expression (green). (**c**) CNO-induced increases in sBG activity as % of pre-CNO baseline, points are 1 unit from 1 bird each. (**d**) Schematic and representative histology for intra-sBG cannula (blue = Neurotrace, red = H3-mCherry, dashed lines = cannula). (**e**) Daily change (Δ) in syllable maturity (last 50 syllables - first 50) 1 day before and during intra-sBG infusion of CNO (paired t-test: t_8_ = 2.44, p = 0.0407). (**f**) No difference in CNO effects on Maturity between intra-sBG and intramuscular (I.M.) CNO, Sal = Saline. (**g**) CNO-induced change (CNO - Saline) in the proportion of canonical syllable transitions (e.g., A→B), no effects detected. (**h**) Acoustic feature means during CNO relative to preceding saline treatment. 2-way ANOVA revealed a main effect of acoustic Feature (F_12,780_ = 2.61, p = 0.0020) and a Feature x Group interaction (int.) (F_12,780_ = 1.85, p = 0.0373). No individual features significantly differed between groups in post hoc tests. (**i**) Change in distance to tutor (au = arbitrary units) from the final saline pretreatment day to 3 days post CNO. No mean difference, but significantly different variance (F_16,12_ = 3.79, p = 0.024). Solid line = median. (**j**) Coefficient of Variation (CV) of acoustic features during CNO relative to preceding saline treatment. 2-way ANOVA revealed a main effect only of group (Ctl/H3, F_1,780_ = 15.7, p < 0.0001). Data points represent individual syllables unless otherwise specified. AM = Amplitude Modulation. meanFreq = mean Frequency. FM = Frequency Modulation. Var = Variance. Scale bars = 500 µm. * = p < 0.05, **** = p < 0.0001. ns = not significant.

## Notes

### Competing Interest Statement

The authors have declared no competing interest.

